# Constitutively active TrkB kinase signalling reduces actin filopodia dynamics and cell migration

**DOI:** 10.1101/2020.09.11.292565

**Authors:** Rohini Gupta, Melanie Bauer, Gisela Wohlleben, Vanessa Luzak, Vanessa Wegat, Dennis Segebarth, Elena Bady, Georg Langlhofer, Britta Wachter, Steven Havlicek, Patrick Lüningschrör, Carmen Villmann, Bülent Polat, Camelia M. Monoranu, Jochen Kuper, Robert Blum

**Affiliations:** Institute of Clinical Neurobiology, University Hospital Würzburg, Würzburg, Germany; Department of Radiation Oncology, University of Würzburg, Würzburg, Germany; Department of Neuropathology, Institute of Pathology, University of Würzburg, Würzburg, Germany; Rudolf Virchow Center for Experimental Biomedicine, Institute for Structural Biology, University of Würzburg, Würzburg, Germany; Comprehensive Anxiety Center, University of Würzburg, Würzburg, Germany; Ludwig-Maximilians-Universität München, Biomedizinisches Zentrum, Planegg, Germany; Fraunhofer-Institut für Grenzflächen- und Bioverfahrenstechnik IGB, Bio-, Elektro- und Chemokatalyse BioCat, Straubing, Germany; Institut für Pathologie, UKE, Hamburg, Germany; Rudolf-Boehm-Institut für Pharmakologie und Toxikologie, Universität Leipzig, Medizinische Fakultät, Leipzig, Germany; Neurona Therapeutics, 170 Harbor Way, South San Francisco, CA, USA

**Keywords:** receptor tyrosine kinase, phosphorylation, NTRK, TrkB, actin, cell migration, FAK, glioblastoma

## Abstract

Trk receptors and gene fusions of *NTRK* are targets in precision oncology. Classical Trk signalling concepts fail to explain ligand-independent signalling of intracellular TrkB or *NTRK* fusion proteins. Here, we show that abundance of the intracellular domain of TrkB is sufficient for ligand-independent autophosphorylation. This constitutive TrkB signalling reduced actin filopodia dynamics, could phosphorylate FAK, and changed cell morphology. Mutating Y^705^ in the kinase domain of TrkB alone specifically blocked these pathways. Engineered intracellular kinase domain proteins and a cancer-related intracellular *NTRK2*-fusion protein (*SQSTM1-NTRK2*) also underwent constitutive activation. In migrating glioblastoma-like U87MG cells, self-active TrkB kinase reduced cell migration. Moreover, we found evidences for constitutively active, intracellular TrkB in tissue of human grade IV glioblastoma. Structural modelling of the kinase domain let us postulate that ‘release from cis-autoinhibition by abundance’ is sufficient for TrkB/FAK/Actin signalling via Y^705^. These constitutive signalling pathways could be fully blocked within minutes by clinically approved, anti-tumorigenic Trk inhibitors. In conclusion, our data provide an explanation and biological function for TrkB kinase domain signalling in the absence of a ligand.

## Introduction

Trk (tropomyosin receptor kinase) receptors belong to the family of membrane bound receptor tyrosine kinases (Barbacid, 1994; Klein *et al*, 1991a; Klein *et al*, 1989; Martin-Zanca *et al*, 1986). The three NTRK genes *Ntrk1*, *Ntrk2*, and *Ntrk3* encode the neurotrophin receptors TrkA, TrkB, and TrkC, respectively (Barbacid, 1994; Klein *et al.*, 1991a). The Trk receptor was originally discovered as oncogenic driver in a human colon carcinoma (Martin-Zanca *et al.*, 1986). Trk receptors or pathological NTRK gene fusions behave pro-tumorigenic in various adult and paediatric tumour types (Cocco *et al*, 2018; Cook *et al*, 2017; Douma *et al*, 2004; Martin-Zanca *et al.*, 1986). Furthermore, overexpression and genomic high-level amplification of NTRK has been observed in a variety of human cancers and abundance of NTRK transcripts can correlate with poor outcomes in affected individuals (Cocco *et al.*, 2018; Eggert *et al*, 2001). To target Trk activity in cancer, membrane-permeable small molecule inhibitors were developed (Roskoski, 2020). Trk kinase inhibitors such as Larotrectinib or Entrectinib are approved modern pharmaceuticals for precision oncology and have high response rates when used to treat NTRK fusion-positive cancers (Cocco *et al.*, 2018; Doebele *et al*, 2020; Drilon *et al*, 2018).

Under physiological conditions, Trk receptors are activated by neurotrophins (Klein *et al.*, 1991a; Levi-Montalcini, 1987; Levi-Montalcini *et al*, 1954; Thoenen, 1995). Neurotrophins are secretory proteins of about 27 kDa (homodimer) and are high-affinity ligands for Trk receptors (Chao & Ip, 2010; Levi-Montalcini, 1987; Levi-Montalcini *et al.*, 1954; Thoenen, 1995). Binding of neurotrophins to extracellular domains of Trk induces receptor dimerization and subsequent trans-autophosphorylation (Chao, 2003; Huang & Reichardt, 2003; Lemmon & Schlessinger, 2010). In TrkA, the transmembrane domain and the juxtamembrane regions carry specific molecular determinants for forming a dimer interface structure that is essential for neurotrophin-dependent Trk dimerization and kinase activation (Franco *et al*, 2020). Ligand-dependent autophosphorylation of Trk recruits a wide variety of cancer-related signalling pathways including the Shc/Ras/Erk-, Pi3K/Akt-. mTOR-, or PLCγ/Ca^2+^ signalling (Chao, 2003; Cocco *et al.*, 2018; Huang & Reichardt, 2003; Sasi *et al*, 2017). The receptor TrkB can also be transactivated in the absence of neurotrophins (Andreska *et al*, 2020; Lee & Chao, 2001). This has been shown in specific types of neurons or neuronal progenitor cells, where membrane-bound, intracellular TrkB becomes activated by metabotropic adenosine (Lee & Chao, 2001; Rajagopal *et al*, 2004; Wiese *et al*, 2007), dopamine signalling (Iwakura *et al*, 2008), or EGF receptor tyrosine kinase signalling (Puehringer *et al*, 2013). Trk receptors can also become constitutively active at intracellular sites, for instance when glycosylation and maturation of the receptor domain is blocked (Watson *et al*, 1999).

While much is known about the physiological Trk kinase signalling, concepts of how atypically activated receptors affect cellular signalling are not fully conclusive (Cocco *et al.*, 2018; Sasi *et al.*, 2017). For instance, in studies looking at NTRK fusion events in cancer, RNA-seq datasets suggest protumorigenic, kinase-active NTRK2 fusion proteins lacking not only the extracellular-ligand-binding domain, but also the transmembrane and juxtamembrane domains (Cocco *et al.*, 2018; Gatalica *et al*, 2019; Stransky *et al*, 2014). This raises the question how intracellular TrkB kinase domains become kinase active, especially when the signals for physiological Trk dimerization are missing (Cocco *et al.*, 2018).

Here we hypothesised that constitutive activation of TrkB at intracellular sites (Chao, 2003; Cocco *et al.*, 2018; Watson *et al.*, 1999) is best explained by self-activation of the intracellular domain (ICD). In our experiments, we confirm that the sole ICD is sufficient for intracellular TrkB kinase activation. Furthermore, we found new signalling properties of TrkB that occur when a high abundance of the kinase domain is available at intracellular sites. In this signalling process, Y^705^ of TrkB, previously known to be involved in kinase activity, is now shown to be directly involved in downstream signalling to cytoskeletal features. We suggest that this TrkB signalling property could be most relevant in case of constitutive activation of TrkB in glioblastoma or NTRK-fusion activity in other types of cancer.

## Results

In order to characterize the functional prerequisite of TrkB for constitutive activation, we used site-directed mutagenesis and PCR techniques to clone a set of mouse TrkB mutants (based on reference sequence: NP001020245, Fig EV1). For overview, Fig 1A shows a model of the TrkB receptor that indicates critical structural components, phosphorylation sites and the binding sites of the antibodies against TrkB used in this study. The specificity and properties of the antibodies were verified with the help of the designed TrkB mutants by means of immunofluorescence (EV2A–D) or Western blotting techniques and are summarized in detail in table EV2E.

**Figure 1.**
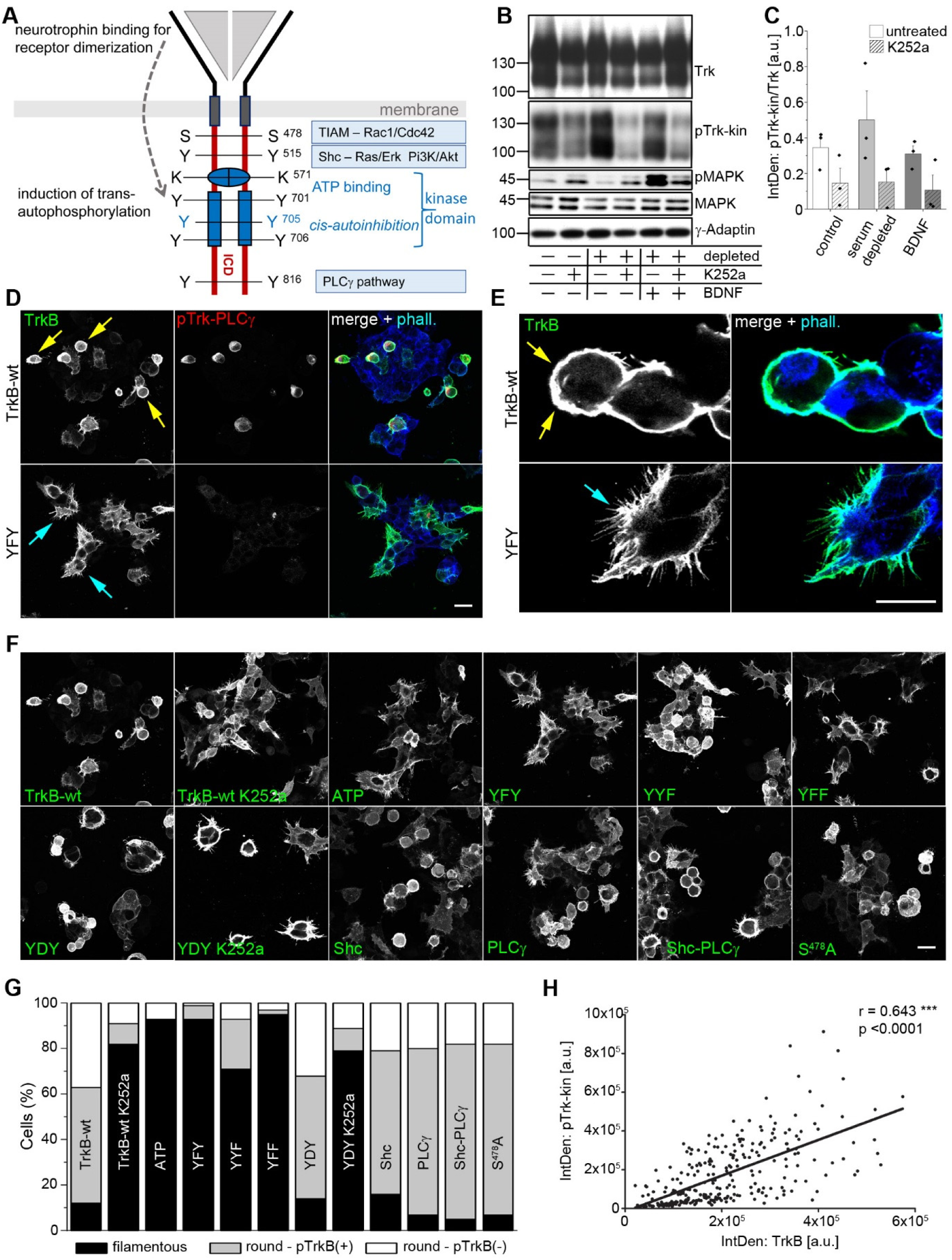
Abundance-dependent self-activation of TrkB kinase and changes in actin morphology strongly depend on phosphorylation of Y705 in the YxxxYY-motif. A Model depicting TrkB-kinase signalling. Neurotrophin (BDNF / NT-4)) binding induces conformational changes and supports receptor dimerization. The kinase is released from cis-autoinhibition, ATP binds to the intracellular kinase domain and allows trans-autophosphorylation. Three tyrosine residues in the consensus motif YxxxYY, Y^701^, Y^705^ and Y^706^ are phosphorylated in the activation loop of the receptor. ATP-binding and phosphorylation of the YxxxYY motif are upstream of further autophosphorylation and downstream signalling. In the intracellular domain (ICD), phosphorylation of the Y^515^ recruits Shc and activates the Ras/ERK and Pi3K-Akt pathways. Y^816^ forms the adaptor site for PLCγ. S^478^ signals to the TIAM-Rac1 pathway. B TrkB phosphorylation in absence of BDNF is unaffected by serum depletion. Western blotting of whole-cell lysates generated from HEK293 cells expressing TrkB. Control cultures were kept in serum before total lysates were produced. Cell cultures were treated as indicated. Serum-depletion was performed for 3h. To inhibit TrkB kinase activity, cultures were preincubated with 150 nM K252a, a Trk kinase inhibitor, for 30 min. DMSO served as solvent control. When indicated, cells were also treated with 10 nM BDNF for 15 min and were compared with BDNF stimulated cells under K252a treatment. C Quantification of Western blots for pTrkB-kin normalized to total TrkB levels with densitometry. Relative integrated densities are shown. K252a treatment for 30 min causes a reduction in TrkB phosphorylation levels under control, serum-depleted and BDNF-stimulation conditions. Bar graph: mean ± SEM, overlaid with single data points; n = 3. D In the absence of neurotrophins, TrkB overexpression induces TrkB phosphorylation and changes in cell morphology. Immunofluorescence of TrkB receptor (green) and pTrk-PLCγ (red). F-actin was labelled with Acti-stain-670 phalloidin (blue). HEK293 cells were transfected with either TrkB-wildtype or TrkB-YFY kinase mutant. Cells were immunostained after 30h. Yellow arrows point to roundish, pTrk-positive cells. Cyan arrows point to filamentous, pTrk-negative cells. Confocal images; scale bar: 25 μm. E Filopodia phenotype of TrkB expressing cells (high-resolution confocal stack image). F Round-shaped cells express kinase-active TrkB mutants. Immunostaining of HEK293 cells expressing either TrkB-wt or indicated TrkB mutants (for details see EV1). Cells expressing TrkB-wt or the constitutive active mutant TrkB-YDY were also subjected to K252a treatment. Typically, filamentous cells express the kinase-dead ATP mutant of TrkB or the YxxxYY mutants (YFY and YFF). Treatment with 150 nM K252a for 30 min reverses the round shape of cells expressing TrkB-wt or TrkB-YDY. Confocal images; scale bar: 25 μm. G Quantification of the percentage of cells showing either a round shape or typical filopodia. Round cells were further subdivided into those that were either positive (+) or not (−) for pTrk. Stainings with anti-pTrk and Acti-stain-670 phalloidin in corresponding TrkB mutants are given in EV3. Data acquired from 10 fields of view in 3 independent experiments. Scale bar: 25 μm. H TrkB phosphorylation correlates with TrkB abundance. Linear, positive correlation of the integrated density of α–TrkB and α–pTrkB-kin immunoreactivity in TrkB-expressing HEK293 cells. Immunolabels per cells were measured as integrated density per cell from maximum intensity projection images of confocal z-stacks. Shown are single cell data, n = 280 cells; data collected from 20 confocal image fields and 4 cell cultures.

### TrkB kinase activity causes abundance-dependent changes in cell morphology

We transiently transfected HEK293 cells with wildtype mouse TrkB (TrkB-wt) and performed Western blotting to test for TrkB expression. TrkB was highly expressed and constitutively active, as expected (Dewitt *et al*, 2014; Watson *et al.*, 1999) (lane 1 in Fig. 1B). Immunoblotting of pTrk was performed with anti-pTrk-kin, an antibody specific for phosphorylation at Y^705^ in TrkB (EV2E). Total and pTrk appeared at 130 kDa, representing the mature, glycosylated TrkB kinase, and 90 kDa, representing a typical immature Trk (Watson *et al.*, 1999). To exclude that serum components in the growth medium were responsible for the pronounced phosphorylation of TrkB, in the absence of neurotrophins, we performed a 3h serum-depletion, which did not reduce the phosphorylation at Y^705^ of TrkB (Fig 1B, C). In addition, we treated the cells with the small molecule K252a, a prototypical and potent Trk inhibitor (Tapley *et al*, 1992). Treatment of HEK293-TrkB cells with 150 nM K252a for 30 min could acutely reduce TrkB phosphorylation, both under control conditions and after 3h serum-depletion (Fig 1B, C). Stimulation with 20 ng/ml BDNF was unable to increase the pTrkB-kin levels any further after serum depletion (Fig 1B, C). This shows that during TrkB overexpression, K252a interrupts an ongoing kinase activity in the absence of neurotrophins. Serum depletion was not sufficient to stop the constitutive activity of overexpressed TrkB, but MAPK phosphorylation downstream of TrkB was reduced (Fig. 1B).

After an expression time of 30–48 h, in the absence of neurotrophins, cells were triple-labelled for TrkB, phospho-TrkB-PLCγ and filamentous actin (F-actin). F-actin was labelled with a phalloidin-Cy5 conjugate, which also stained the gross morphology of the cell. The majority of TrkB-wt expressing cells had a round cell body and showed a pronounced Trk phosphorylation signal (Fig 1D, G). In contrast, the Y^705^-TrkB mutant (TrkB-YFY) did not show an obvious phosphorylation at the PLCγ-site (pPLCγ), indicating a reduction of autophosphorylation in this mutant (Fig 1D). Furthermore, TrkB-YFY expressing cells formed typical filopodia (Fig 1E). This indicates that Y^705^ is important for regulating cytoskeletal features in constitutively active TrkB.

Next, we determined the potency of diverse TrkB mutants to affect the F-actin in HEK293 cells (Fig 1F, G, EV3). Transfection of cells with the kinase-dead TrkB mutant (TrkB-ATP) or mutants of the kinase Y^705^ (TrkB-YFY and TrkB-YFF) did not alter actin/TrkB+ filopodia formation and the cells showed, like untransfected HEK293 cells, typical filopodia (EV3). However, cells expressing either TrkB-wt or any of the other tested TrkB mutants for the Shc, PLCγ or TIAM/Rac1/CDC42 interaction sites showed a roundish cell shape. In TrkB-wt cells and in cells expressing the constitutive active TrkB (TrkB-YDY), the roundish cell phenotype could be reversed by K252a treatment for 30 min, indicating that kinase activity is involved in the observed cell morphology changes. For downstream signalling of TrkB, the Shc- and PLCγ-adapter sites are crucial (Chao, 2003; Huang & Reichardt, 2003). However, even the double mutant TrkB-Shc-PLCγ disrupted the filopodia-like phenotype (Fig 1F, G, EV3), indicating that the classical Shc/PLCγ signalling pathways of TrkB did not cause the change in cell morphology.

To test for abundance effects, we overexpressed TrkB in HEK293 cells and labelled them with either anti-TrkB or anti-pTrkB-kin. We determined the integrated density of corresponding immunolabels on the single cell level in confocal microscope images. The data confirmed a linear, statistically significant correlation between the abundance of TrkB and pTrkB-kin intensity (Fig 1H).

TrkB protein is expressed as either the TrkB receptor kinase or as the kinase-deficient TrkB splice isoform TrkB-T1 (Klein *et al*, 1991b; Middlemas *et al*, 1991). TrkB-T1 consists of the complete extracellular region and transmembrane domain but carries only a short cytoplasmic tail of 23 amino acids. Overexpression of TrkB-T1 (Rose *et al*, 2003) did not cause a round cell morphology and did not destroy filopodia formation (EV4A,B). Overexpression of the other Trk kinase family members, TrkA and TrkC, also caused a roundish cell shape (EV4C,D). We did not look further at TrkA and TrkC, but instead focused solely on TrkB for the scope of this study.

Confocal imaging of the TrkB phosphorylation signal suggested that most of the phospho-Trk-wt signal was localized at intracellular sites (Fig 1D). For this reason, we performed a series of control experiments to test whether constitutively active TrkB is localized at intracellular sites, as observed earlier for TrkA (Watson *et al.*, 1999). These experiments revealed that TrkB-wt, but not kinase-dead TrkB-ATP, shows a tendency to be already active at intracellular sites (EV5A). Furthermore, mutating 12 predicted N-glycosylation sites in the receptor domain did not block constitutive TrkB activation (EV5B). Finally, TrkB cell surface labelling confirmed phospho-active TrkB at intracellular sites, in the absence of a ligand (EV5C).

### Actin filopodia formation is disturbed in TrkB overexpressing cells

To better describe the actin-phenotype in TrkB-expressing cells, confocal live cell imaging was performed (Fig. 2). Cells were co-transfected with TrkB, TrkB mutants (representatively shown in Fig 2 is the mutant TrkB-YFF), and GFP-actin. In TrkB-wt cells, two phenotypes were typically observed: (1.) round cells with low GFP-actin dynamics, or (2.) some single cells forming bleb-like structures (Fig 2A, video EV 6,7). Bleb formation is a type of cell motility that is observed when the cytoskeleton is decoupled from the plasma membrane (Charras *et al*, 2006). In contrast, cells expressing TrkB-YFF showed typical actin filopodia dynamics (Fig. 2B, video EV 8,9).

**Figure 2.**
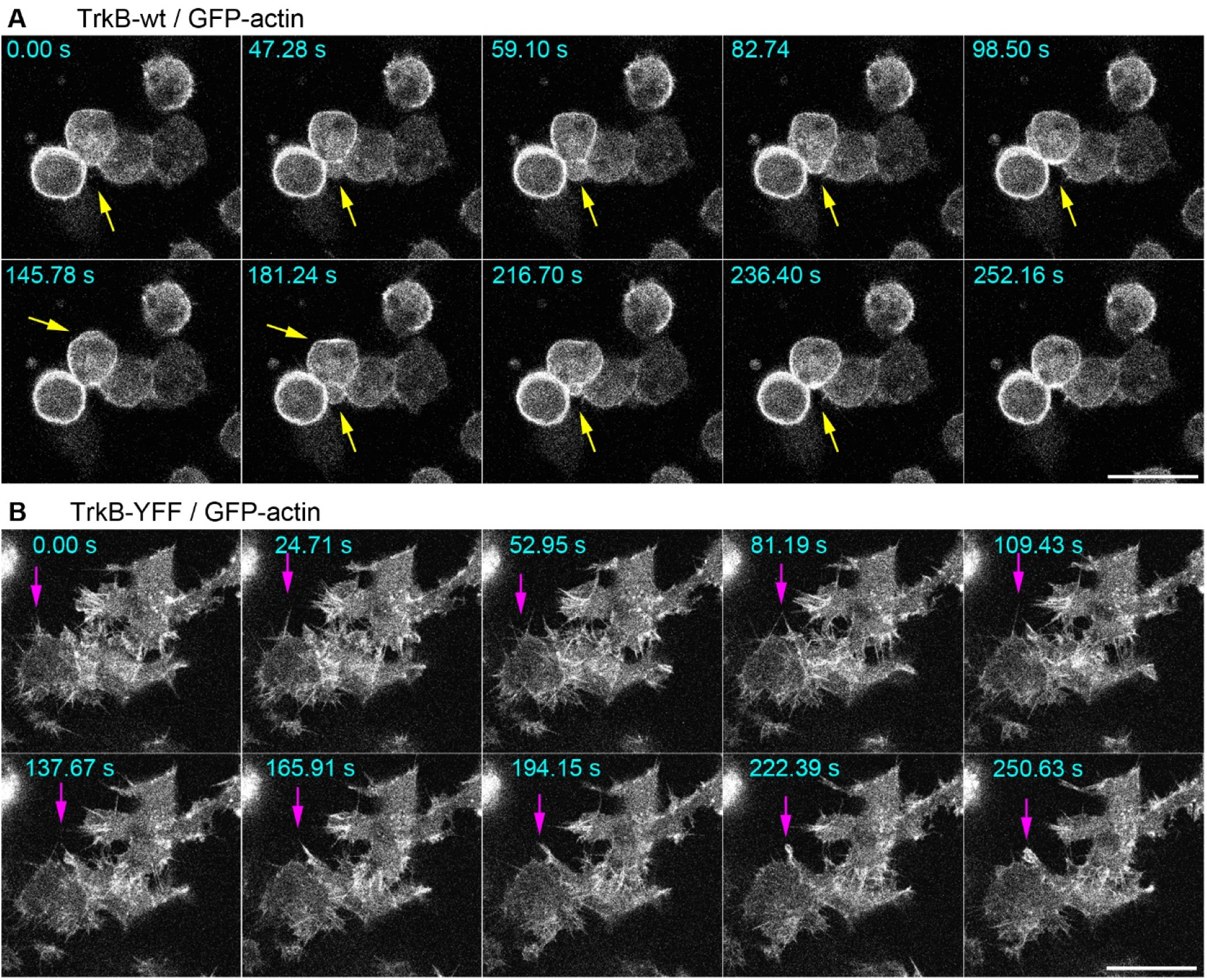
Mutating of the YxxxYY-motif in TrkB restores actin filopodia dynamics. A, B Overexpression of TrkB kinase induces a round-shaped cell morphology and loss of actin filopodia dynamics. HEK293 cells co-expressing GFP-actin and TrkB-wt (in A) or the TrkB-YFF mutant. Time-lapse images (in seconds) are shown. Living cells were imaged using a confocal laser-scanning microscope. GFP-actin was excited with a 488 nm laser line and fluorescence was detected with a spectral detector (510 – 570 nm). (In A) Arrows point to a roundish cell that forms typical blebs. (in B) Arrow points to typical dynamic filopodia, labelled by GFP-actin. The corresponding time-lapse videos are provided as extended videos (EV6, TrkB-wt, overview; EV7 TrkB-wt, blebbing; EV8 Trk-YFF overview; EV9 TrkB-YFF, filopodia). Scale bar: 25 μm.

### TrkB overexpression induces phosphorylation of Focal Adhesion Kinase (FAK)

The blebs observed in actin live cell imaging led us to ask whether constitutively active TrkB kinase induces the phosphorylation of proteins involved in actin dynamics (Blanchoin *et al*, 2014; Parsons *et al*, 2010). We overexpressed diverse TrkB mutants and probed total cellular protein with anti-Trk and anti-pTrk (PLCγ). Furthermore, we tested for phosphorylated Cofilin, a protein involved in reorganization of F-actin, and Focal Adhesion Kinase (FAK), a cytosolic tyrosine kinase regulating focal adhesion site assembly, membrane protrusion formation and cell motility (Parsons *et al.*, 2010). Anti-phospho-Cofilin immunoblotting was inconspicuous, but a strong phosphorylation of FAK at Y576/577 was seen in TrkB-wt expressing cells (Fig 3A). FAK phosphorylation could be acutely inhibited by the Trk inhibitor K252a (Fig. 3B,D). Cells expressing the kinase-dead TrkB-ATP mutant, or cells expressing TrkB with a site-directed missense mutation in Y^705^ or Y^706^ did not show phosphorylation of FAK. Moreover, constitutively active TrkB-YDY, a protein typically associated with a round cell phenotype (Fig. 1G,F), did not show pFAK activation (Fig. 3A). This indicates that the round cell phenotype depends on kinase-active TrkB which can be mimicked by substituting Y^705^ with D^705^ or E^705^ but this does not necessarily cause pFAK activation. The data show that pFAK activation by TrkB depends on Y^705^ and Y^706^.

**Figure 3.**
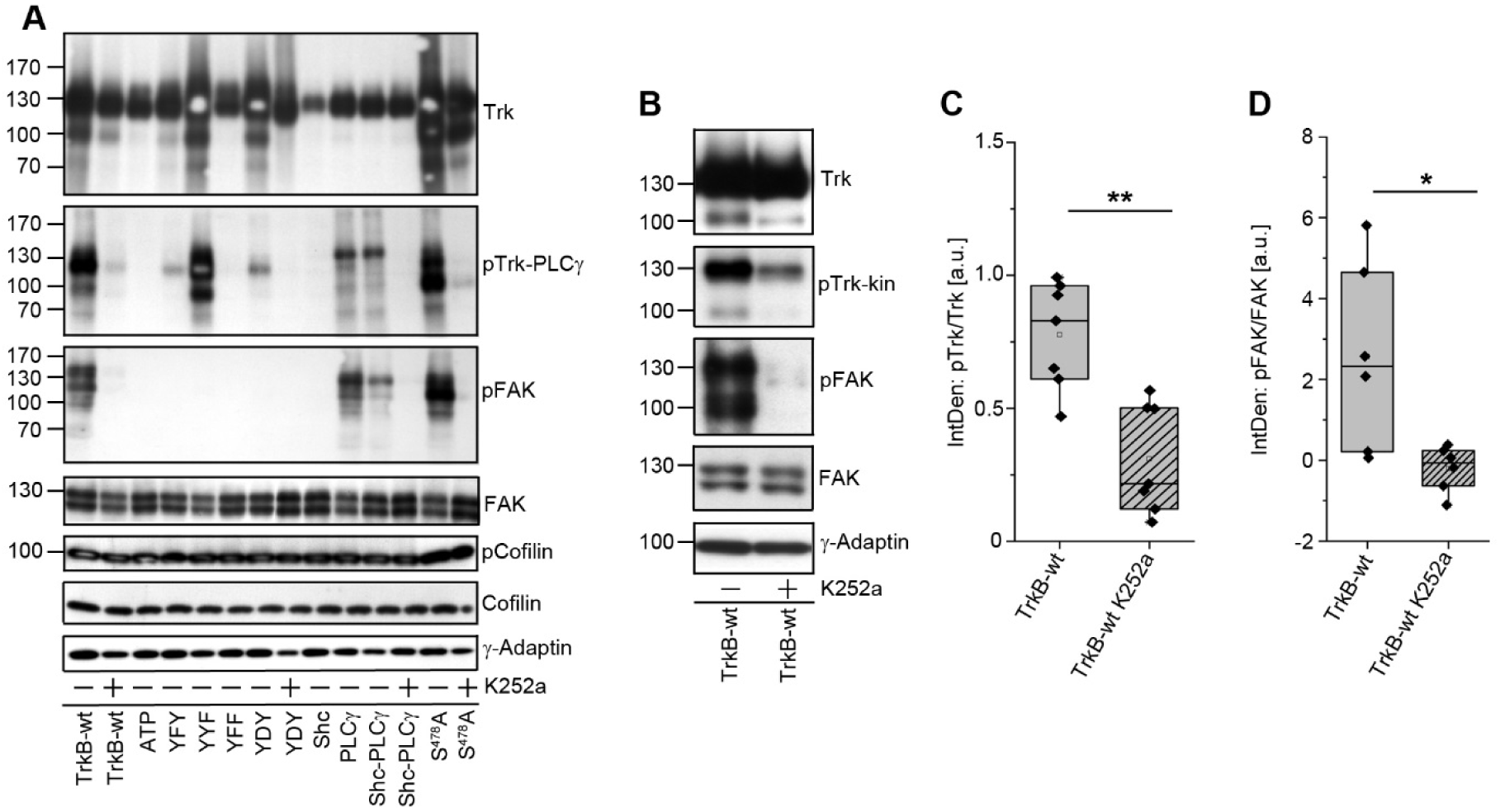
Self-active TrkB is upstream of Focal Adhesion Kinase (FAK) phosphorylation. A Self-active TrkB with an intact kinase domain induces phosphorylation of Focal Adhesion Kinase (FAK), a key player of actin dynamics, but not of Cofilin. Western blotting of whole-cell lysates generated from HEK293 cells expressing indicated TrkB mutants. To inhibit TrkB kinase activity, cultures were preincubated with 150 nM K252a for 30 min. DMSO served as solvent control. Antibodies against total FAK, Cofilin and γ-Adaptin served as loading control. B Self-active TrkB induced phosphorylation of FAK can acutely be blocked with the Trk kinase inhibitor K252a. Western blotting of whole-cell lysates generated from HEK293 cells expressing TrkB-wt. To inhibit TrkB kinase activity, cultures were preincubated with 150 nM K252a for 30 min. DMSO served as solvent control. C Quantification of Western blots for pTrkB-kin normalized to total TrkB levels with densitometry. Relative integrated densities are shown. K252a treatment causes a reduction in TrkB phosphorylation levels. Bar graph: mean ± SEM, overlaid with single data points; n = 7; 2-sample-t-test was performed: t(12) = 4.295, p = 0.00104. D Quantification of Western blots for pFAK normalized to total FAK levels with densitometry. Relative integrated densities are shown. K252a treatment causes a reduction in FAK phosphorylation levels. Bar graph: mean ± SEM, overlaid with single data points; n = 6; Mann-Whitney-U-test was performed: U=32, p = 0.03064.

### The intracellular domain of TrkB is sufficient to induce FAK phosphorylation

Most of the constitutively active pTrkB was found at intracellular sites (EV5). As FAK is a cytosolic protein kinase, we asked whether cytosolic expression of the intracellular domain of TrkB would be sufficient to induce FAK phosphorylation. To test this, we cloned two prototypical intracellular kinase domain constructs. The construct TrkB-ICD, carried the complete intracellular domain of TrkB (K^454^ to C-terminal end) (Fig 4A). Myr-ICD, consisted of the TrkB ICD coupled to an aminoterminal myristoylation (Myr) / S-acylation targeting motif, combined with a GGSGG-linker sequence. This motif was used to target Myr-ICD to the plasma membrane (Kabouridis *et al*, 1997; Rathod *et al*, 2012) (Fig 4A). We expressed both constructs in HEK293 cells, performed immunolocalization experiments (Fig. 4B), and probed total protein lysates with diverse antibodies (Fig. 4C). Immunolabelling confirmed that both, TrkB-ICD and Myr-ICD were phosphoactive at Y^705^/Y^706^, albeit the cellular localization profile was different (Fig 4B). TrkB-ICD appeared at intracellular sites throughout the cytosol (Fig 4B, magenta arrows), while phospho-active Myr-ICD outlined the cell surface, indicating its efficient targeting to the plasma membrane from the intracellular site (Fig 4B, yellow arrows). Western analysis confirmed that ICD-protein and Myr-ICD were phospho-active and migrated at the predicted relative molecular weight of 40-45 kDa. Surprisingly, the ICD domain was sufficient to cause a dramatic induction of pFAK phosphorylation at Y^397^ (Fig 4C, E) and at Y^576/577^. The constitutive active ICD did not induce MAPK phosphorylation (Fig 4C, F). In striking contrast, Myr-ICD caused almost no FAK phosphorylation, but led to activation of MAPK (Fig. 4C). Again, TrkB-wt was able to induce both, pFAK and pMAPK, and K252a could reduce pTrk and pFAK signals (Fig 4B-F). The kinase mutant TrkB-YFF did not show pFAK activation, but induced MAPK phosphorylation (Fig 4F). From these data we assume that Myr-ICD, when targeted to the plasma membrane, can provide a platform for adapter proteins and thereby support a certain level of constitutive Ras/MAPK signalling, even in the absence of neurotrophins. The fact that the ICD alone does not promote MAPK activation is in line with earlier data by the group of R.S. Segal showing that unglycosylated, constitutively active TrkA, in tunicamycin-treated PC12 cells, does not activate MAPK (Erk) (Watson *et al.*, 1999).

**Figure 4.**
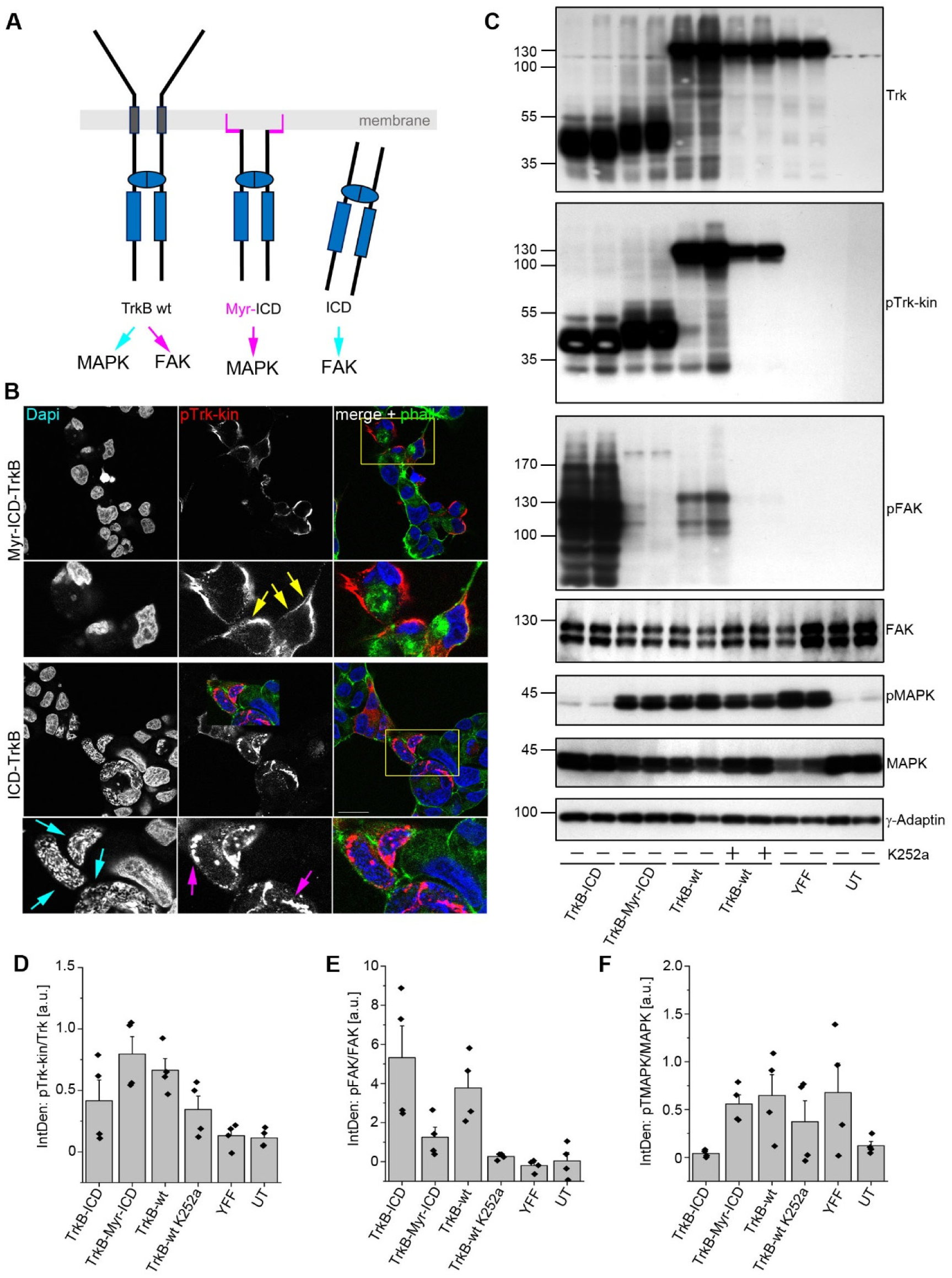
The intracellular domain (ICD) of TrkB transduces signals to FAK, while a membrane-targeted ICD of TrkB is linked to MAPK phosphorylation. A Intracellular, kinase-active domain constructs of TrkB. In Myr-ICD, an N-terminal myristoylation consensus motif with a glycine-serine (GGSGG)-linker was used to target the ICD to the plasma membrane. In contrast to TrkB-wt, the ICD (intra<u>c</u>ellular <u>d</u>omain) construct lacks the ligand-binding and transmembrane domain. The model includes the result of the experiment. Constitutive active TrkB-wt signals to MAPK and FAK, while the Myr-ICD activates MAPK, but not FAK. The ICD activates FAK, but not MAPK. B TrkB-ICD and TrkB-Myr-ICD undergo self-activation but differ in their cellular localization pattern. Immunofluorescence of pTrk-kin (red) and DAPI (blue). F-actin was labelled with Acti-stain-670 phalloidin (phall, green). HEK293 cells were immunostained 30 h after transfection. TrkB-ICD is preferentially seen at intracellular sites. The Myr-ICD construct is shows typical plasma membrane targeting. Note morphology changes in DAPI of ICD-expressing cells, indicating changes in chromatin compaction (cyan arrows). Single plane confocal images; scale bar: 20 μm. C TrkB-ICD, but not TrkB-Myr-ICD induce FAK phosphorylation. Western blotting of whole-cell lysates generated from HEK293 cells expressing TrkB-ICD, TrkB-Myr-ICD, TrkB-wt and TrkB-YFF. K252a was used to inhibit Trk kinase activity. TrkB-wt expression increases MAPK and FAK phosphorylation. ICD signals to FAK, but not MAPK. In striking contrast, Myr-ICD induces MAPK phosphorylation, but fails to activate FAK. D-F Quantification of Western blots for pTrkB-kin normalized to total TrkB levels (in D), pFAK to total FAK (in E) and pMAPK to total MAPK (in F). Relative integrated densities are shown. Bar graph: mean ± SEM, overlaid with single data points; n = 4.

### Structural features of TrkB with phosphorylated or unphosphorylated Y^705^

Our data show that Y^705^ in TrkB plays an important role in abundance-dependent activation and downstream signalling of TrkB. Structural models for the activation mechanism of TrkB are mainly based on generalized models of receptor tyrosine kinase activation (Artim *et al*, 2012; Hubbard *et al*, 1994). These models suggest that in the absence of a ligand, receptor tyrosine kinases are autoinhibited in cis and that autoinhibition is released following ligand-induced receptor dimerization. In order to understand the molecular basis of this activation, we had a closer look at Y^705^ utilizing a TrkB crystal structure (pdb code 4AT4). If self-activation and constitutive signalling depends mostly on Y^705^, we asked whether there are unusual structural features in TrkB compared to TrkB-YFY that allow for intracellular self-activation in the absence of a ligand. We used the TrkB model for molecular dynamics (MD, GROMACS) and performed a 1 ns run. The resulting structure was compared to the kinase domain of the insulin receptor (1gag) to see how a fully triple phosphorylated autoinhibition loop looks like (Fig 5A). We observed large structural rearrangements between the TrkB model and the activated insulin receptor. Interestingly the inactive form of the insulin receptor closely matched the conformation of the MD TrkB model, indicating a similar autoinhibited state (Fig 5A). In a next step, we performed identical MD runs using two TrkB variants (phospho Y^705^ and Y^705^F) and compared the resulting models. *In silico* phosphorylation of Y^705^ in the TrkB model suggests that the phosphorylation changes the autoinhibition loop position. This is indicated by the significant shift of the YxxxYY motif (3.7 Å at Gly^712^ located at the tip of the loop, Fig 5B). Interestingly a similar change can be observed with the Y^705^F variant. These small but significant differences indicate structural transitions in the receptor structure that may underlie TrkB activation by a ligand-independent release from cis-autoinhibition upon overexpression.

**Figure 5.**
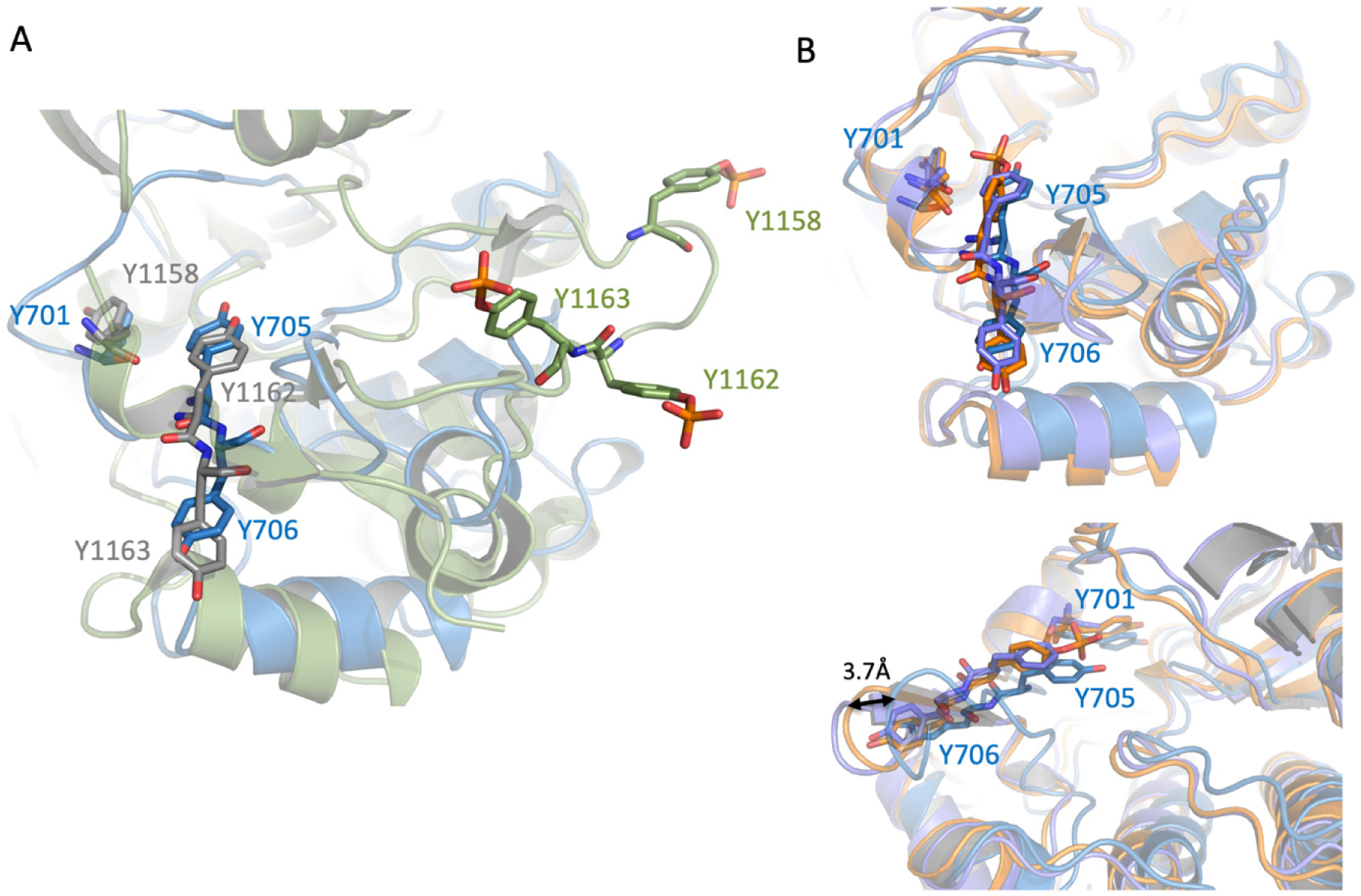
Modelling of TrkB. (A) Superimposition of TrkB MD model (blue) with the activated form of the insulin receptor (1gag, green). The YxxxYY motif is depicted in ball and stick mode. The YxxxYY motif of the autoinhibited insulin receptor is shown in grey and ball and stick mode. The backbone has been omitted for clarity. (B) The upper panel shows a superimposition of three MD models for TrkB. Wild type is shown in blue. Phospho Y^705^ in orange and Y^705^F in purple. The YxxxYY motif is shown in ball and stick mode and the movement is indicated by an arrow.

### Trk inhibitors acutely block constitutive Trk and cytosolic NTRK2-fusion signalling

The clinically approved Trk inhibitors LOXO-101 (Larotrectinib) and Entrectinib are used to suppress classical and oncogenic Trk activity (Doebele *et al.*, 2020; Drilon *et al.*, 2018), but whether they would also block downstream signalling of intracellular TrkB has not been tested. Here we show that all small molecule Trk inhibitors efficiently blocked constitutive TrkB activation and pFAK phosphorylation within 60 min (Fig. 6A,B). Constitutive pMAPK activation by TrkB (see Fig. 4C,F) was efficiently blocked by LOXO-101 and Entrectinib (Fig. 6). A side observation was that K252a could block TrkB kinase self-activation and pFAK downstream of Trk. However, in case of K252a, MAPK remained active (see Fig. 1C or Fig. 6B,D, lane 2). We looked at this effect in more detail and saw that 150 nM K252a promotes phosphorylation of MAPK, for instance in serum-depleted cells (Fig. 1B) and potentiates EGF signalling to MAPK (EV10). This shows an unexpected off-target effect of K252a.

**Figure 6.**
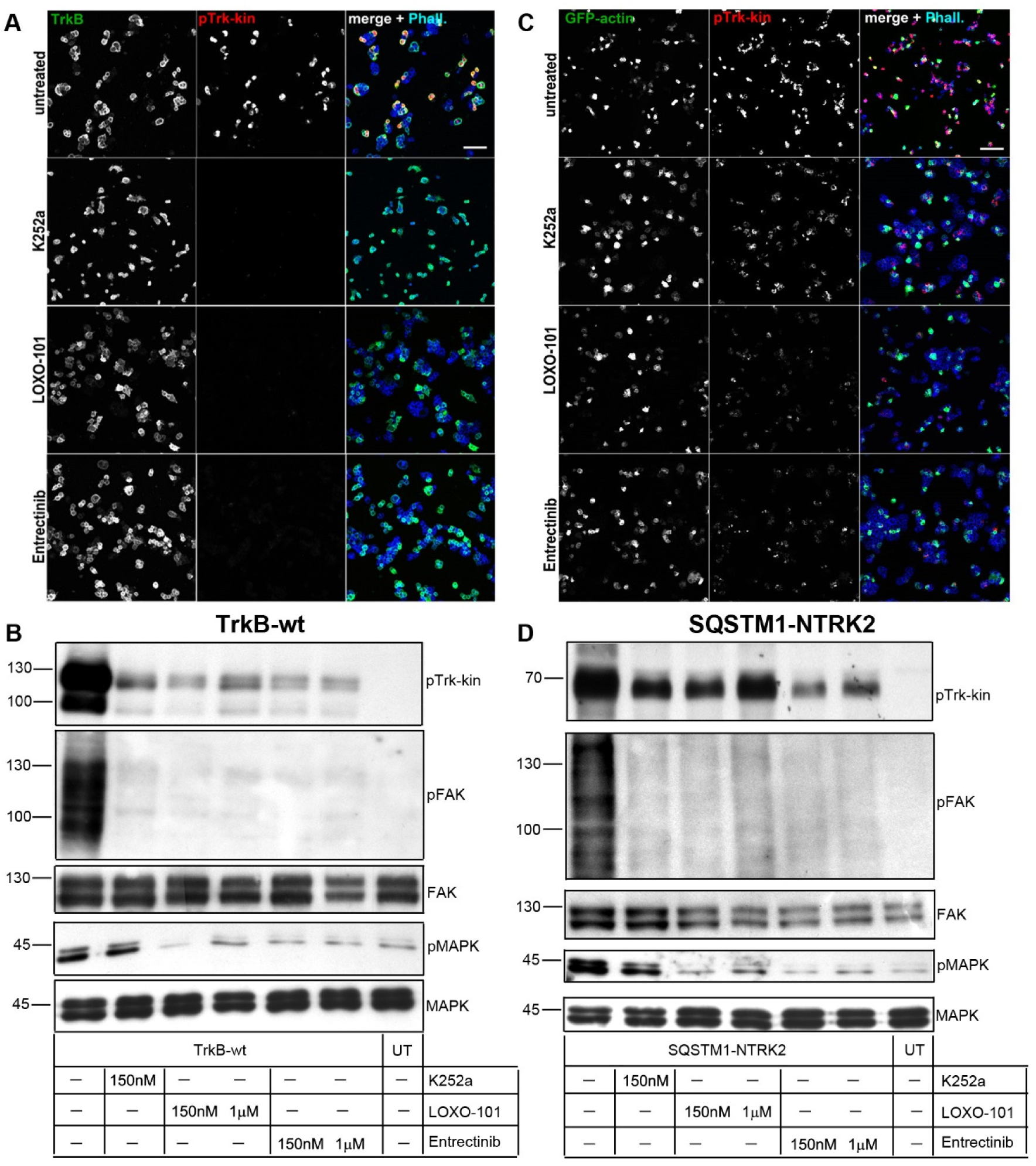
Treatment of TrkB-wt and SQSTM1-NTRK2 (a TrkB fusion product) with small molecule Trk inhibitors reduces or hinders downstream Trk activated pathways. A Immunostaining of HEK293 cells expressing TrkB-wt. Immunofluorescence of TrkB (green), pTrk-kin (red) and Acti-stain-670 phalloidin (phall, blue). Cells were treated with small molecule Trk inhibitors like K252a, LOXO-101 and Entrectinib. TrkB self-activity is reduced upon inhibitor treatment. Confocal images; scale bar: 100 μm. B Western blotting of whole-cell lysates generated from HEK293 cells expressing TrkB-wt. After transient transfection, TrkB was expressed for 30 h and then treated with small molecule Trk inhibitors like K252a, LOXO-101 and Entrectinib (concentrations are as depicted). Lysates were probed with the indicated antibodies. Trk, FAK and MAPK activity is reduced upon inhibitor treatment. C Immunostaining of HEK293 cells expressing a synthetically designed TrkB fusion construct – SQSTM1-NTRK2. This particular fusion product was chosen because it lacks the TrkB juxtamembrane domain and the Shc site, thereby making it primarily an intracellular protein with an intact kinase domain. Cells were co-transfected with GFP-actin. Immunofluorescence of GFP-actin (green), pTrk-kin (red) and Acti-stain-670 phalloidin (phall, blue). Cells were treated with small molecule Trk inhibitors like K252a, LOXO-101 and Entrectinib. TrkB fusion self-activity is reduced upon inhibitor treatment. Confocal images; scale bar: 100 μm. D Western blotting of whole-cell lysates generated from HEK293 cells expressing TrkB fusion construct – SQSTM1-NTRK2. After transient transfection, TrkB fusion was expressed for 30 h and then treated with small molecule Trk inhibitors like K252a, LOXO-101 and Entrectinib (concentrations are as depicted). Lysates were probed with the indicated antibodies. While Trk activity is not hampered, downstream activity of FAK and MAPK is reduced upon inhibitor treatment.

Some cancer-related NTRK fusion proteins lack the aminoterminal receptor domain and are fused to cytosolic proteins (Martin-Zanca *et al.*, 1986; Stransky *et al.*, 2014). Structural features suggest that these proteins are localized at intracellular sites and become oncogenic drivers due to kinase activation by a yet elusive mechanism (Cocco *et al.*, 2018). In order to investigate whether such NTRK-fusion proteins behave like the TrkB-ICD, we expressed the protein SQSTM1-NTRK2 in HEK293 cells. SQSTM1-NTRK2 was found in a lower grade glioma in RNA-seq data. In this fusion protein, exon 1 – 5 of sequestosome 1 (SQSTM1), a multifunctional signalling adapter involved in autophagy, are fused to exon 16 - 20 of NTRK2 (Cocco *et al.*, 2018; Gatalica *et al.*, 2019; Stransky *et al.*, 2014). This creates an open reading frame and links the aminoterminal part of SQSTM1 with the kinase domain of human TrkB. The TrkB domain includes the complete kinase region and an intact C-terminus but lacks the Shc adapter site and the juxtamembrane region. SQSTM1-NTRK2 showed constitutive phosphorylation, caused a roundish cell phenotype, was able to induce phosphorylation of FAK^Y576/577^ and, in contrast to the Trk-ICD, also MAPK (Fig. 6C,D). The Trk kinase inhibitors K252a, LOXO-101 and Entrectinib blocked the kinase-related, constitutive activation of FAK and MAPK.

Together, these data indicate intracellular signalling of constitutively active TrkB to actin filopodia dynamics and FAK which are both involved in cell migration. Therefore, we used migratory-active glioblastoma-like cells to investigate constitutive active TrkB signalling in cell migration.

### Constitutive active TrkB inhibits migration of U87MG cells

Human U87MG glioblastoma-like cells are commonly used in brain cancer research but have an unknown patient origin. The clone U87MG (ATCC) is of CNS origin and carries bona fide glioblastoma-like characteristics (Allen *et al*, 2016). The cells are migratory and are suited to better understand how candidate proteins interfere with non-directed cell migration (Diao *et al*, 2019).

To better control protein abundance in this cell model, we expressed the TrkB-wt, TrkB-YFY, and the NTRK gene fusion construct SQSTM1-NTRK2 in a doxycycline-inducible lentiviral expression system (Wang *et al*, 2014). A second resistance gene cassette was used to select Trk-positive cells with 1μg/ml puromycin after lentiviral transduction. In absence of doxycycline, expression of all proteins was below detection limits (exemplarily shown for TrkB-wt in Fig. 7 and EV11). However, induction of expression with 1μg / ml doxycycline for 48 h led to TrkB expression and constitutive activation of TrkB (Fig. 7B, EV11). In this cell system, TrkB was strongly enriched close to F-actin-rich protrusions (EV11, arrows in magenta). Notably, two fractions of pTrk-kin immunoreactivity were observed. The most prominent pTrk-kin signal was close to the perinuclear region, in a Golgi apparatus-like localization. This signal was negative for the receptor domain antibody (EV11, cyan arrows; see also EV5). The distribution pattern of TrkB and F-actin was reminiscent of typical cell morphology in non-migrating cells (Etienne-Manneville, 2008). The intracellular protein SQSTM1-NTRK2 also became constitutively active (Fig. 7B, EV11).

**Figure 7.**
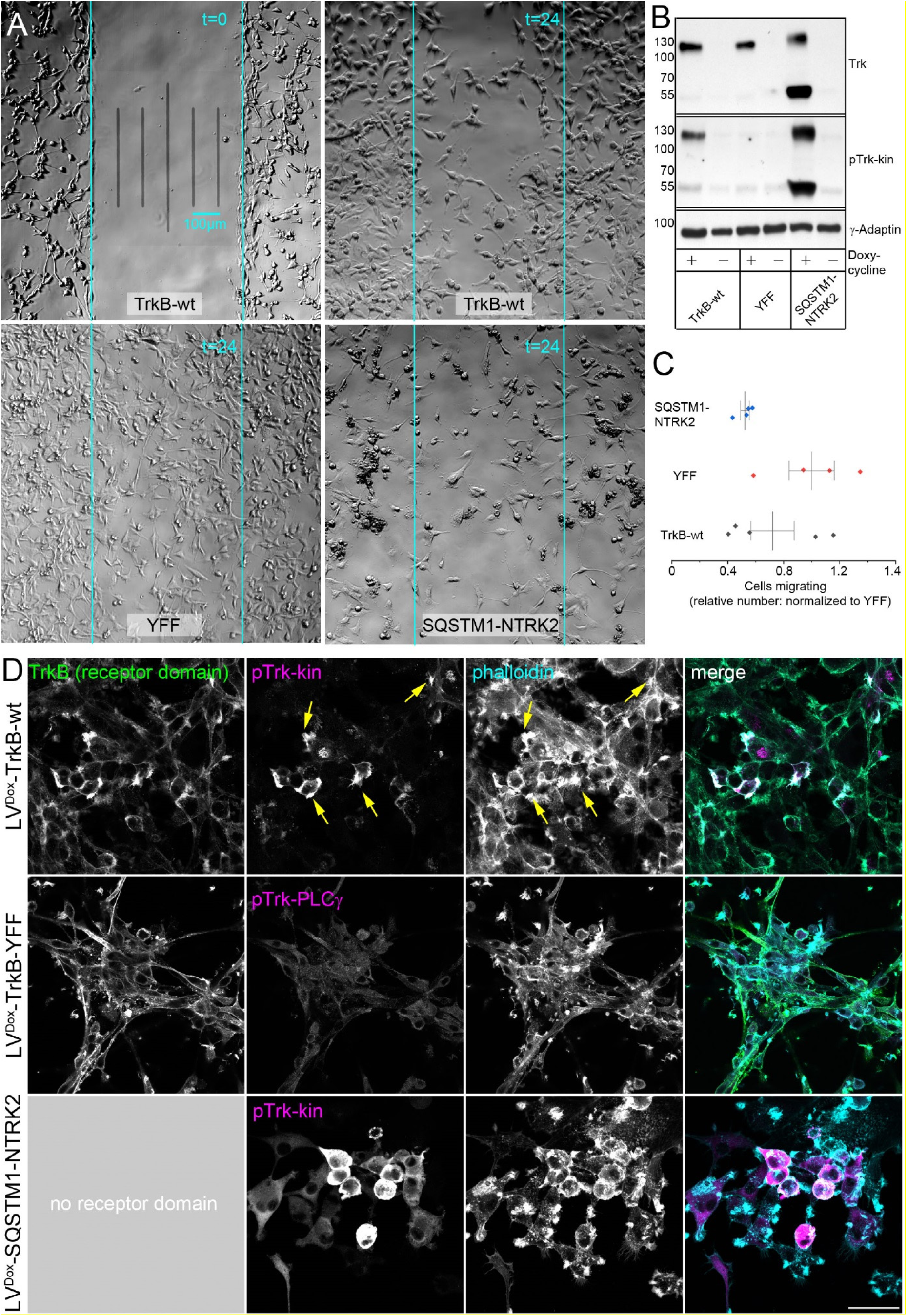
Constitutive active TrkB kinase reduces the migratory activity of glioblastoma-like cells. A Representative phase contrast microscopy images of migration assay. Shown are U87MG cells expressing either TrkB-wt, TrkB-YFF or the fusion protein SQSTM1-NTRK2. The image in the upper left corner shows the cells in culture after removing the silicon insert (t=0). The other images representatively show the situation 24 h later. Note the cell morphology changes in TrkB-wt and SQSTM1-NTRK2 expressing cells versus TrkB-YFF. B Western blotting of whole-cell lysates generated from U87MG cells expressing indicated TrkB kinase construct. Cells transduced with indicated lentiviral constructs were kept as polyclonal cell line and puromycin was used to select transduced cells. In absence of Doxycycline, cells did not express the corresponding proteins. Doxycycline was added to induce Trk-construct expression for 48h. Constitutive activation of Trk was verified with pTrk-kin. γ–Adaptin = loading control. C Migratory activity of U87MG cells, expressing indicated Trk-kinase constructs. Shown are relative numbers of cells to the initially cell-free gap. Cell count was normalized to the mean of TrkB-YFF expressing cells. Migratory activity is shown relative to TrkB-YFF, which expresses the same structural protein domains as TrkB-wt, but is mutated at Y^705^ and Y^706^. (See also EV12) D Immunostaining of U87MG cells expressing inducible TrkB-wt, TrkB-YFF and the NTRK fusion construct SQSTM1-NTRK2. Immunofluorescence of TrkB receptor domain (green), pTrk-kin (red) and Acti-stain-670 phalloidin (phall, blue). Yellow arrows point to constitutive pTrk close to F-actin. TrkB-YFF-expressing cells were labelled with anti-pTrk-PLCγ because the antibody binding site of anti-pTrk-kin is mutated in this construct (EV2E). SQSTM1-NTRK2 does not have a receptor domain, as indicated. Confocal z-stack images; scale bar: 50 μm.

Next, we seeded the U87MG cells expressing the different constructs into 2-well silicon inserts with a defined cell-free gap for testing random migration (Fig. 7A). With cell seeding, we added 1 μg/ml doxycycline to induce the expression of the three constructs. For control, we added the solvent DMSO. After another 24 h, we removed the silicon insert and observed whether induction of TrkB did speed up or slow down random cell migration. The experiment showed that expression induction significantly reduced the cell migration of TrkB-wt expressing cells compared to TrkB-YFF expressing cells (Fig. 7A, C). Morphological alterations and cell migration effects of SQSTM1-NTRK2 expressing cells were extreme (Fig. 7A,D, EV11). The cells were roundish. Cell migration was rather weak (Fig. 7A,C), but cell clone formation was observed (Fig. 7A, EV11). In absence of neurotrophins, the pure abundance and expression of the construct was responsible for the dramatic changes of the cellular properties with respect to random migration and cell clone formation. Another interesting observation was that the SQSTM1-NTRK2 protein, with a predicted molecular weight of 61 kDa appeared at about 60 and 120 kDa under standard SDS-PAGE Western blotting conditions (Fig. 7B). This indicates that the protein tends to aggregate and to form a rather stable, SDS-resistant dimer. After expression induction, U87MG cells expressing TrkB-wt, TrkB-YFF or SQSTM1-NTRK2 retained their ability to grow (EV12).

Up to here, our experiments revealed an unusual, novel signalling property of the intracellular Trk kinase domain via Y^705^. However, Trk effects were seen in a rather artificial situation, namely constitutive Trk signalling after recombinant expression of TrkB or TrkB mutants. Consequently, we examined human glioblastoma (grade IV) tissue samples, to investigate, whether we find evidence for intracellular, constitutively active TrkB under pathological, yet physiological conditions.

### TrkB kinase in human glioblastoma samples

The TrkB-encoding gene NTRK2 is abundantly expressed in neural cells during neural development and in the adult brain. Besides that, the receptor is also highly abundant in diverse types of glioblastoma, the most common tumours of the brain (Wadhwa *et al*, 2003). Constitutive Trk receptor signalling is pro-tumorigenic in glioblastoma (Lawn *et al*, 2015; Wang *et al*, 2018) and fusions of NTRK1, NTRK2, and NTRK3 genes belong to the genomic landscape of diverse types of gliomas (Cook *et al.*, 2017; Lawn *et al.*, 2015; Wu *et al*, 2014). In this context, we asked whether there is evidence for intracellular Trk-kinase signalling in grade IV glioblastoma.

Glioblastoma tissue was harvested during brain surgery of patients suffering from glioblastoma (first diagnosis or recurrence, male and female, age 33 – 80 years old). For this study, we took tissue categorized by histological examination as grade IV glioma. Frontal brain tissue (post-mortem, male and female, 33 – 72 years old) was used as control. To localize pTrk in glioblastoma tissue, we took frozen sections and performed immunofluorescence labelling with anti-Nestin. High Nestin expression can be used to distinguish glioma cells from unaffected brain tissue with high probability (Ma *et al*, 2008; Zhang *et al*, 2008). Morphologies ranged from Nestin-positive clone-like cell clusters, cells rich in neurites, to cell clumps or dense cell masses (EV13). Cryosections were then labelled with anti-Nestin and anti-TrkB. We could identify TrkB in many Nestin+ cells in certain areas of the glioma tissue (EV14). Then we labelled for pTrk and Nestin and performed high-resolution z-stack confocal microscopy of pTrk-positive cells (Fig. 8A). Single Nestin+ cells carried intracellular pTrk-positive clusters and even pTrk-positive bleb-like formations (Fig. 8A). It is important to note that the observation was seen in single cells or selected areas within the rather heterogeneous glioblastoma multiforme. To be sure that the tissue expresses the TrkB kinase splice variant, we harvested tissue RNA from cryosections, isolated the RNA and performed reverse-transcriptase qPCR (Fig. 8B). The upper primer was positioned to an exon encoding for the receptor domain and the lower primer was bound to an exon encoding for the TrkB kinase domain (table 3). By this strategy, transcripts for truncated TrkB-T1 are not detected. Relative expression was compared to RNA-Polymerase II transcripts and revealed a rather high expression level of the TrkB kinase transcript in the control samples of frontal brain tissue (about 50% of the housekeeping gene RNA polymerase II, Fig. 8B). In glioblastoma cryosections a rather high, but variable expression of TrkB kinase was found (Fig. 8B).

**Figure 8.**
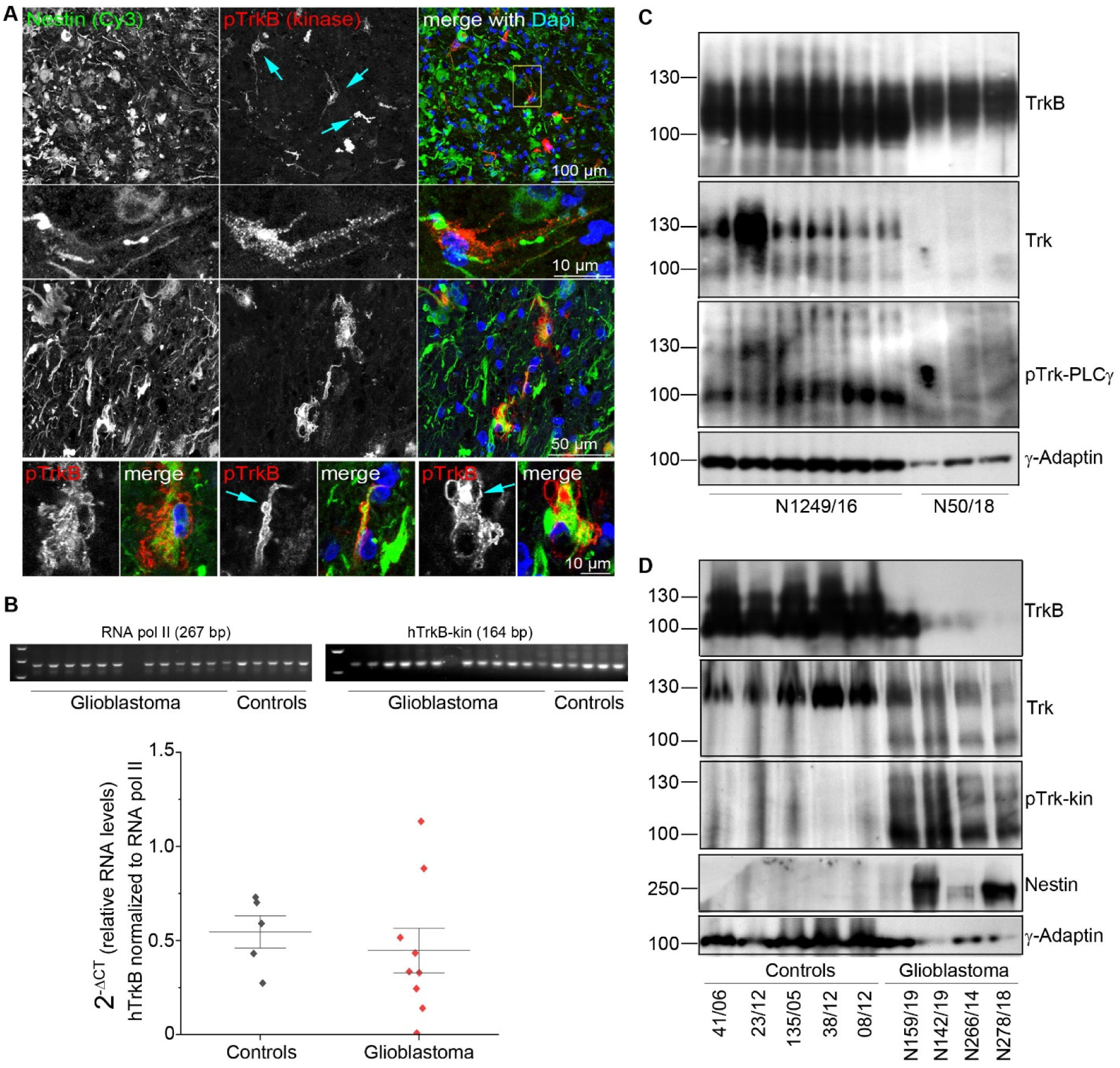
Localization and expression of TrkB, Trk and pTrk in grade IV glioblastoma. A Immunofluorescence analysis of a representative glioblastoma tissue cryosection. Confocal images are shown. Sections were labelled for anti-Nestin, for identification of the glioblastoma and pTrk. Single Nestin+ cells show a high abundance of pTrk (upper panel, cyan arrows). Second and third panel: High-resolution confocal image stack showing pTrk in intracellular, immunoreactive clusters. Lower panel. Membrane bleb-like structures in single cells with strong intracellular pTrk label. B RT-qPCR reveals the abundant expression of the TrkB kinase transcript in grade IV glioblastoma. TrkB kinase expression levels are given in relation to RNA polymerase II. Single data points, the mean and the standard deviation are indicated. The size of the amplicons was verified by agarose gel electrophoreses, as indicated. C TrkB and phospho-active Trk kinase in glioblastoma. Western blotting of whole-cell lysates generated from frozen, post-mortem glioblastoma samples with indicated antibodies. Lane 1–6 shows lysates generated from different tissue pieces of the same glioblastoma sample. Note high abundance of Trk kinase and pTrk in one tissue piece (lane 2). Lane 7-9 shows a representative TrkB-positve, Trk kinase negative, pTrk-negative glioblastoma sample. γ-Adaptin, loading control. D TrkB and phospho-active Trk kinase in glioblastoma. Western blotting of whole-cell lysates generated from human brain samples or glioblastoma samples. Lane 1–5 shows lysates generated from control brain samples (frontal brain). Trk kinase immunoreactivity is seen at 130 kDa, indicating mature Trk. Lane 6 – 9 represent total lysates from different grade IV glioblastoma. Note high abundance of Trk kinase and pTrk at about 90 kDa, indicating immature, phosphorylated Trk.

Next, we took bigger tissue samples (approximately 1cm^3^), dissected these in smaller pieces and prepared total cell lysates. Fig. 8C shows representative Western blots for two samples, a sample from a first diagnosis (N1249/16) and a recurrence sample from another patient (N50/18). Due to the neural origin of glioblastoma, probing samples with anti-TrkB gave a pronounced signal around 90 – 100 kDa, indicating the rather strong expression of the truncated TrkB-T1. Due to the strong immunoblotting signal at 90 kDa, we could not clearly resolve the mature TrkB-kinase at 130 kDa. Probing with anti-Trk, an antibody detecting all Trk isoforms, verified the expression of Trk-kinase (Fig 8C). The band at 90 KDa may suggest expression of the non-glycosylated immature Trk as well as the typical 130 kDa band. Notably, the Trk kinase receptor was not equally distributed in the tissue. One of the six tissue pieces showed a much higher abundance of Trk kinase (Fig 8C, lane 2). Furthermore, probing with pTrk (PLCγ-site)-antibodies confirmed strong phosphorylation signal at 90 kDa, suggesting phosphorylation of the immature Trk isoform (Fig 8C). In one of the lysates with very high Trk kinase protein expression, the 130 kDa band also showed an anti-pTrk signal (Fig 8C). In the second tissue sample (N50/18), neither the Trk kinase, nor the pTrk could be detected (Fig. 8C). Next, we compared protein lysates from human frontal brain with protein lysates from different grade IV glioblastoma samples (recurrence samples) (Fig. 8D). Again, we saw pronounced TrkB and Trk-kinase signals in Western blots. However, the 90 kDa Trk kinase signal was exclusively observed in glioblastoma samples. Trk kinase in frontal brain controls showed the typical 130 kDa mature Trk band (Fig 8D). pTrk signals were abundant in glioblastoma and barely visible in control brain tissue (Fig 8D). This experiment points to immature, phospho-positive Trk kinase in glioblastoma and shows that this is a common phenomenon.

## Discussion

Like all tyrosine kinase receptors, the TrkB receptor is activated by dimerization and subsequent autophosphorylation of intracellular tyrosine residues (Jura *et al*, 2011; Lemmon & Schlessinger, 2010). Release of cis-autoinhibition, following ligand-induced receptor dimerization, is the key event that has been proposed to trigger receptor tyrosine kinase activation (Artim *et al.*, 2012; Bertrand *et al*, 2012; Hubbard *et al.*, 1994; Lemmon & Schlessinger, 2010). However, classical concepts for ligand-dependent TrkB kinase activation fail to fully explain signalling mechanisms of intracellular TrkB (Watson *et al.*, 1999), or cancer-related, cytosolic NTRK fusion proteins (Cocco *et al.*, 2018).

### Trk activation and release from cis-autoinhibition

Our data are in accordance with observations in the 90s of the last century, showing that the isolated ICD of TrkB undergoes abundance-dependent autophosphorylation (Iwasaki *et al*, 1997). This biphasic effect was thought to be mediated by a intramolecular (cis) activating step and a second abundance-dependent intermolecular (trans) step (Iwasaki *et al.*, 1997). In our Myr-ICD or ICD constructs, typical dimerization domains such as the juxtamembrane region or the single span transmembrane domain (Franco *et al.*, 2020) are missing. We assume that the release of cis-autoinhibition happens in ICD monomers and subsequently a high abundance of ICD domains is needed to enable transactivation between ICDs. In this concept, cis-autoinhibition is not a stable conformation. Sequential cis/trans-activation and subsequent downstream signalling might be a contributing factor for ligand-independent TrkB functions. *In silico* phosphorylation of Y^705^ or the Y^705^F mutation, cause a structural shift (Fig. 5). This transition could be indicative for the concept that the release from cis-autoinhibition initiates Y^705^-mediated signalling. Why phosphorylation and a variant (Y^705^F) that cannot be phosphorylated have a similar outcome remains, however, elusive.

### Self-activation of TrkB ICDs – an option for constitutive signalling by Y^705^

Downstream signalling of TrkB typically occurs via the TIAM/Rac -, Shc-adapter -, or PLCγ/Ca^2+^-dependent signalling cascades (Cocco *et al.*, 2018; Huang & Reichardt, 2003; Sasi *et al.*, 2017). The finding that Y^705^ is critical for intracellular, constitutive activation of TrkB is, in the context of the literature, at least at a first view, not surprising (Chao, 2003; Cocco *et al.*, 2018; Franco *et al.*, 2020; Iwasaki *et al.*, 1997; Watson *et al.*, 1999). What’s surprising is rather the following: (1.) Persistent activity of intracellular Y^705^ of TrkB is upstream of a very specific signalling event that interrupts actin filopodia dynamics. (2.) It can inhibit random cell migration and (3.) induce focal adhesion kinase (FAK) phosphorylation. (4.) It persists in the absence of neurotrophins or serum components and (5.) can be stopped acutely with Trk inhibitors.

The signalling is different from typical BDNF-dependent (Klein *et al.*, 1991b) or neurotrophin-independent TrkB activation by adenosine, EGF or dopamine (Iwakura *et al.*, 2008; Lee & Chao, 2001; Puehringer *et al.*, 2013; Rajagopal & Chao, 2006; Rajagopal *et al.*, 2004; Wiese *et al.*, 2007). Intracellular ICD-mediated, constitutive activation is an independent category of signalling options of the TrkB kinase domain via Y^705^. We thus suggest referring to this phenomenon as ‘TrkB kinase self-activation’.

### TrkB ICD-signalling to FAK

Constitutive activation of TrkB interrupts actin filopodia formation. Cells become round and show membrane blebbing (Fig. 1, Fig. 2). This led us to investigate whether focal adhesion kinase (FAK) (Westhoff *et al*, 2004), is downstream of Y^705^ signals. However, we saw that Y^705^ signalling to FAK and to cell migration are different events. Constitutive active TrkB-wt, but also YxxxYY-mutants such as YYF, YDY, or YEY, all interrupt actin filopodia formation, but do not activate FAK phosphorylation (Fig. 2, Fig. 3 and Fig.4). Under controlled expression, TrkB also undergoes self-activation and interrupts cell migration. We consider it important, because it might be part of a yet undefined NTRK-fusion signalling pathway or ligand-independent TrkB signalling in cancer. The specificity of Y^705^ signalling is remarkable because it is blocked in so-called ‘constitutive active YDY’ mutants, or YYF mutants. In future studies, it will be interesting to find out whether a high on-off dynamic of the pathway might contribute to the turnover rate of focal adhesion sites in TrkB expressing cells in absence of ligands, before transactivating ligands or BDNF promote actin filopodia dynamics and chemotactic migration events.

### Constitutive TrkB kinase self-activation in grade IV glioblastoma?

In human grade IV glioblastoma tissue, we found marked differences in the TrkB profile, when compared with frontal brain control tissue. In GBM, Trk kinase phosphorylation was strong in the 90 kDa TrkB band, typical for non-glycosylated, immature TrkB. Immunolocalization and high-resolution microscopy confirmed intracellular pTrk-positive clusters, at least in some Nestin+ cells. These results and do not exclude the possibility that the natural TrkB ligand BDNF from neurons, microglia, serum (Naegelin *et al*, 2018) or platelets (Fujimura *et al*, 2002) is stimulating Trk phosphorylation. However, it has been shown that recombinant expression of a TrkB construct lacking the immunoglobulin-like domains of TrkB, meaning the BDNF-binding domain, is sufficient to confer an aggressive carcinogenic phenotype to a neural crest-derived cell line (Dewitt *et al.*, 2014). Therefore, in the context of the overall literature and our results here, we should be open for the possibility that there is, among other signalling pathways, also abundance-dependent self-activation of endogenous, unmutated intracellular TrkB, which is upstream of cytoskeletal functions and pro-tumorigenic. Because of this idea, we suggest testing multiple tissue pieces of GBM biopsies (first diagnosed and recurrence samples) for pTrk-Y^705^ abundance. In clinical studies, it should be tested whether anti-Trk treatment can contribute to a better outcome for glioblastoma patients with a pronounced pTrk signal.

Furthermore, our study suggests that a pharmacologic, small molecule block of the release from cis-autoinhibition at Y^705^ or the stabilization of the YFY-like confirmation (Fig. 5) would be rather interesting strategic options to block constitutively active NTRK in cancer, for instance in case of acquired resistance to prior Trk kinase inhibition (Drilon *et al*, 2017; Okamura *et al*, 2018).

## Materials and Methods

### Cloning and plasmids

All mammalian expression vectors used in this study were constructed in pcDNA3, or in the lentiviral vector FuGW (Lois *et al*, 2002). We refer to the cDNA sequence *Ntrk2* (*trk*B full-length – *trk*B.FL) (reference: NM_001025074 / NP001020245) for all mouse TrkB constructs. Doxycycline-inducible lentiviral expression of TrkB constructs was performed with a vector backbone based on pCW (Wang *et al.*, 2014). The pCW vector expresses a puromycin resistance gene cassette under the ubiquitous hPGK1 promoter. All constructs used in this study are listed in table 1. Lentiviral constructs in FuGW carried an aminoterminal HA-tag between a signal peptide and the first amino acid of the mature TrkB receptor (Nikoletopoulou *et al*, 2010). Mutants were generated with site-directed mutagenesis using synthetic oligonucleotides and Quick Change II XL mutagenesis kit (Agilent Technologies). An additional human *Ntrk2* fusion construct (SQSTM1 fused to TrkB kinase (Stransky *et al.*, 2014) was generated by gene synthesis (Eurofins).

**table 1.**
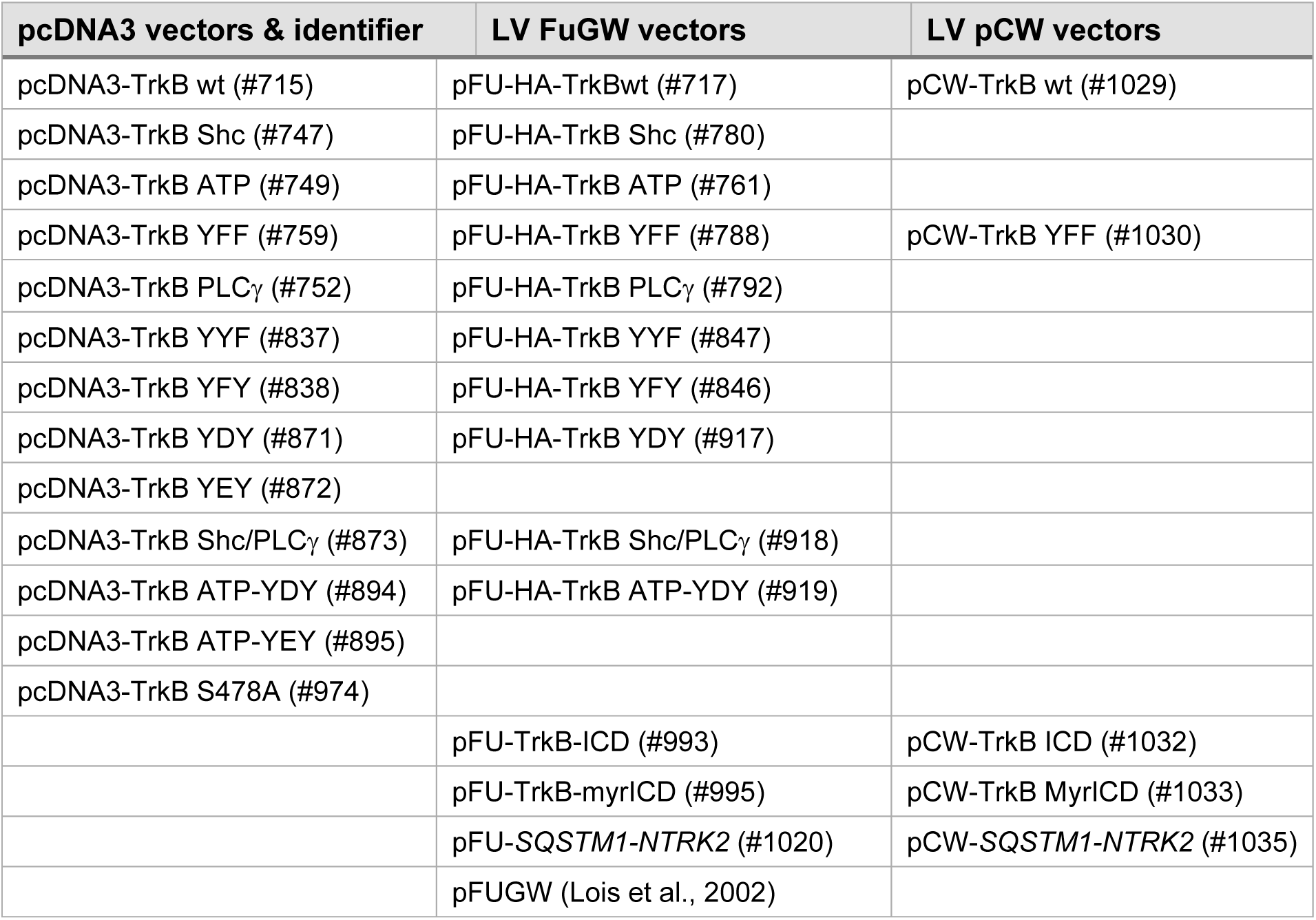
mammalian expression vectors.

### Cell culture and transfections

HEK293 cells (ACC #305) were grown in DMEM, high glucose, with GlutaMAX (Gibco), 10% FCS and 100 units / ml of penicillin and 100 μg / ml of streptomycin (Gibco). U87MG, a glioblastoma-like cell line (Allen *et al.*, 2016), was purchased from ATCC (#HTB-14) and grown in the same medium. Cells were incubated at 37°C in 5% CO_2_. For transfection, lipofectamine 2000 (Invitrogen) was used in a ratio 1 μg DNA per 2 μl lipofectamine. Media was replaced after 24 h and the expression was maintained for 30 h – 72 h.

### Lentivirus and LV-generated TrkB-expressing U87 MG cells

Lentiviral particles were packaged in HEK293TN producer cells (SBI biosciences) with pCMV-VSVG and pCMVΔR8.91 (Zufferey *et al*, 1997) helper plasmids. Cells were transfected with Lipofectamine 2000 (Invitrogen) in OptiMEM medium with 10% FCS for 12–14 h. Viral supernatants were harvested 72 h after transfection by ultracentrifugation. Viral particles were suspended in (in mM) 50 Tris–HCl, pH 7.8, 130 NaCl,10 KCl, 5 MgCl_2_ and stored at −80°C. The viral titer was tested in HEK293 cells. The number of infectious particles was determined using serial dilutions of the viral vectors on HEK293 cells. For transduction of U87MG, a multiplicity of infection (m.o.i.) of one was used. One day after transduction, U87MG cells were cultured in presence of 1 μg/ml puromycin.

### Cell migration assay

U87MG cells expressing the TrkB constructs were seeded at a density of 20.000 cells per well into a 2-well silicone insert (Ibidi, #81176), positioned in a 35 mm μ-dish (35mm, high, Ibidi, #81156). One day after seeding, 1 mg/ml doxycycline was added to induce expression of the corresponding TrkB-related constructs (Dox on). For control, the solvent DMSO was added. 24 h after Dox-induction, the cell culture dishes were filled with growth medium and the silicone insert was removed. Cells were monitored using brightfield microscopy directly after removal of the insert and after 24 h, to analyze cell migration. Subsequently cells were fixed and labelled by immunofluorescence.

The detection and quantification of cells in the acquired brightfield images was automatized with custom written code in ImageJ (Rasband, W.S., ImageJ, U.S. National Institutes of Health, Bethesda, Maryland, USA, https://imagej.nih.gov/ij/). Images were converted to 8-bit grayscale and smoothed by replacing each pixel with the mean of its neighbourhood (5-pixel radius). Cells were defined as local maxima with a prominence greater than 7.5 in the smoothened images (representative images shown in EV12). The area of the removed silicon insert was annotated manually in each image and the number of cells detected within this area was quantified.

### Antibodies

Primary antibodies were used for indirect immunofluorescence labelling or Western blot analysis at indicated dilutions (see table 2).

**table 2.**
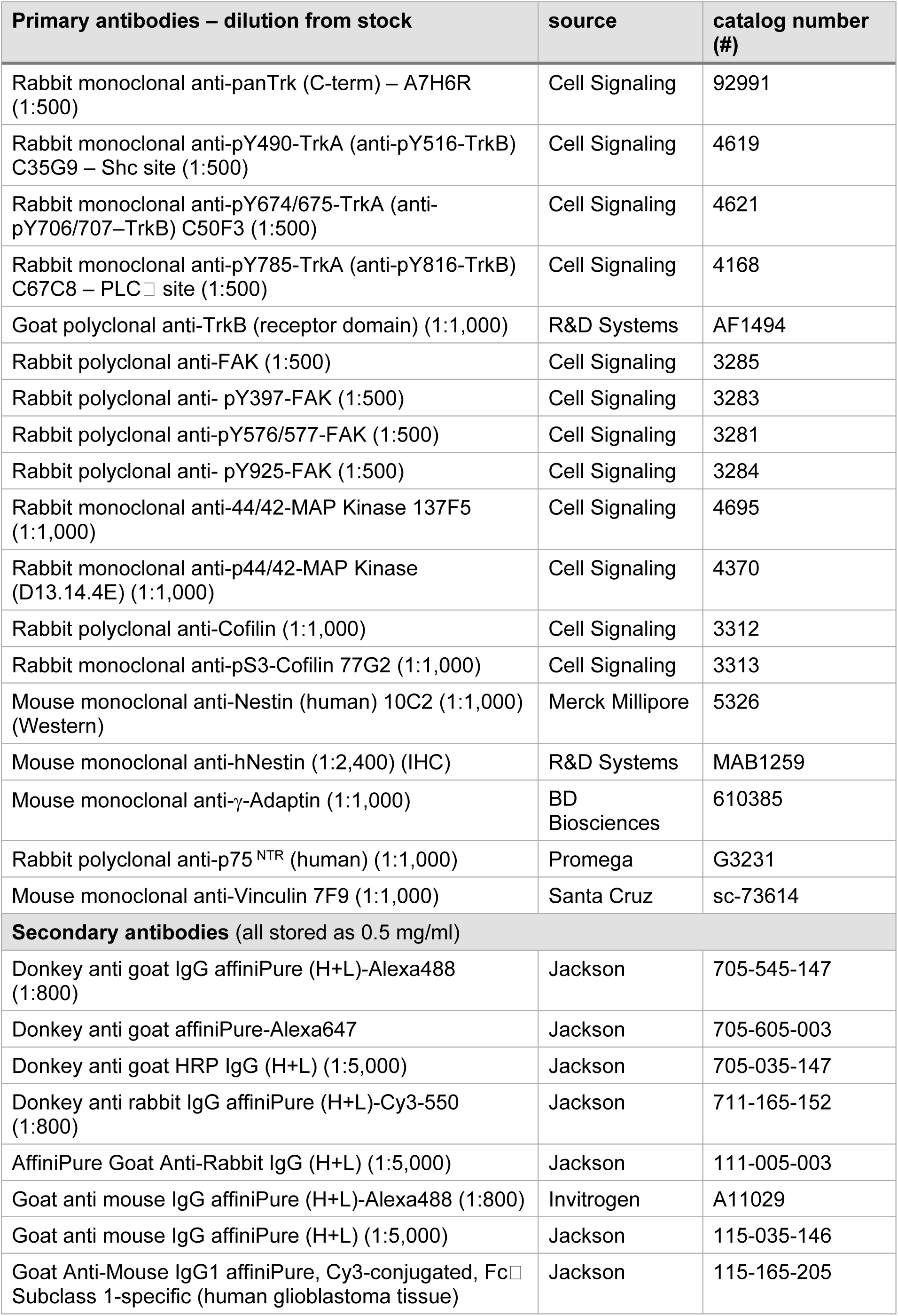
antibodies.

### Indirect immunofluorescence

Coverslips (10 mm, Marienfeld) were placed in 4-well tissue culture dishes (Greiner) and coated with 0.1 mg/ml poly-L-lysine (PLL, Sigma). Cells were seeded on these coverslips (150,000 cells per dish). After transfection and indicated expression times, cells were fixed with PBS-buffered 4% paraformaldehyde (pH 7.4) for 15 min at 37°C. Blocking solution contained 1% BSA or 10% horse serum in PBS supplemented with 0.1% Triton X100 and 0.1% Tween 20. Antibodies were diluted in blocking solution. Coverslips were then washed 8 times in 0.1% Tween 20/PBS). Fluorochrome-conjugated secondary antibodies (Alexa-Fluor 488, Cy3, and Cy5 (Jackson laboratories) were used for 1 h at room temperature (21 – 23°C). Cell nuclei were labelled with DAPI (2 mg/ml stock solution, freshly diluted 1:5000 in PBS) for 5 min at RT. For some experiments, cells were also further incubated for 30 min with Alexa-670-phalloidin (Cytoskeleton #PHDN1) to label for actin filaments. Following DAPI treatment, cells were washed twice with PBS. The coverslips were finally mounted with Aquapolymount (Polyscienes).

### Confocal Laser scanning microscopy and image processing

Images were acquired using an inverted IX81 microscope equipped with an Olympus FV1000 confocal laser scanning system, a FVD10 SPD spectral detector and diode lasers of 405, 473, 559 and 635 nm. All images shown were acquired with an Olympus UAPO 20x (air, numerical aperture 0.70) or UPLSAPO 60x (oil, numerical aperture:1.35) objective. For high-resolution confocal scanning a pinhole setting representing one Airy disc was used. In case of high-resolution imaging, confocal settings were chosen to meet an optimum resolution of at least 3 pixels per feature in x-y direction. In z-direction, 300 nm steps were used. 12-bit z-stack images were processed by maximum intensity projection and were adjusted in brightness and contrast using Image J software (Rasband, W.S., ImageJ, U.S. National Institutes of Health, Bethesda, Maryland, USA, https://imagej.nih.gov/ij/) (Schneider *et al*, 2012). Images are shown as RGB images (8-bit per colour channel). Fluorescence images were processed for final presentation using Adobe Photoshop CS5.

### Live Cell Imaging

For live cell imaging experiments, μ-high 35 mm Ibidi dishes (Ibidi #81156) were utilized. These dishes were first coated with poly-L-ornithine (PORN). HEK293 cells were grown on the coated dishes – 100,000 cells per dish. Cells were transfected with TrkB mutants and GFP-actin plasmids. After 24h, old media was replaced with prewarmed HEPES-buffered DMEM (containing 10% FCS, 100 units / ml of penicillin and 100 μg / ml of streptomycin and 1 mM sodium pyruvate). Cells were imaged using Leica SP5 inverted confocal microscope equipped with Leica objectives (HC PL Apo×20/0.7; HCX Apo ×60/1.4–0.6 oil). GFP actin was excited with a 488 nm laser line. Fluorescence was detected with a spectral detector (12 bit) at Airy disc1 settings.

### TrkB kinase domain modelling

As basis for the modelling process, the pdb entry 4AT4 was chosen. MD runs were performed using the gromacs 5.1 package. After solvation, the addition of ions, energy minimization, and equilibration, productive MD was run for 1 ns. The TrkB variants were generated using the program COOT and simulated with the same protocol as the original model.

### Western blot analysis

HEK293 cells were grown on 35 mm cell culture dishes (Falcon). 200,000 cells per dish were transfected with DNA plasmids using Lipofectamine 2000. Cells were lysed at 30h on ice using a cell scraper and 150 μl cold lysis buffer (1% NP40, 50 mM HEPES pH 7.5, 150 mM NaCl, 10% Glycerol, 1 mM sodium fluoride, 10 mM sodium pyrophosphate, 2 mM sodium orthovanadate, 5 mM EDTA, supplemented with one EDTA-free protease inhibitor mini tablet / 5 ml of buffer (Roche #4693159001). Lysates were incubated on ice for 15 min, sonicated twice for 5 s (Hielscher sonifier UP50, M1 sonotrode, 80% power, 10 cycles à 0.5 s) and placed back on ice for 10 min. Lysates were centrifuged for 5 min at 4°C at 15,000xg. Protein concentration of the samples was determined by Pierce BCA protein assay kit (Thermo Scientific). For SDS-PAGE, the supernatant was mixed with Laemmli sample buffer and heated for 5 min at 95°C. After SDS-PAGE, proteins were transferred onto PVDF membranes (Immun-Blot, BioRad). Western blotting was done at 4°C using Mini Trans-Blot Cell Assembly (BioRad) at 25 V, 0.3 A for 15 h. Blocking solution contained 5% milk powder (BioRad) in Tris-buffered saline containing 0.2% Tween 20. After blocking for 30 min at RT, primary antibody were used in blocking solution for 3 h at RT. Blots were washed thrice (20 min each) with TBST and then incubated with secondary antibody for 2 h at RT. Blots were developed using ECL (Immobilon Western HRP Substrate, Merck Millipore) and X-ray films (Fujifilm Super RX). For quantification, these developed X-ray films were first photographed with a 12MP Canon camera by placing them on a white transilluminator plate inside the PeqLab Gel Documentation system and were saved as 12-bit images. The integrated densities were then calculated for the protein bands using ImageJ.

For analysis of human glioblastoma tissue, frozen tissue samples were briefly thawed at RT and washed twice in 1x PBS. Small chunks were then distributed into microfuge tubes and lysed with cold lysis buffer as mentioned above. Sonification and centrifugation steps were repeated as needed until a homogenous suspension was obtained. Samples were loaded onto SDS-PAGE gels after BCA protein quantification and then immunoblotted, as described above.

### RNA isolation and quantitative RT–PCR

Patient glioblastoma frozen samples were thawed and washed twice in 1x PBS. Small chunks were then distributed into microfuge tubes and RNA isolation was performed using the Rneasy Mini Kit (Qiagen). To generate cDNA, Superscript III Reverse Transcriptase first strand synthesis kit (Invitrogen, #12371-019) was used with 500 ng RNA and 50 ng random hexamer primer. cDNA reaction was 5-times diluted in 10 mM Tris-HCl, pH8.5 containing 1 mg/ml BSA. Light cycler 96 Detection System (Roche) was used to perform RT-qPCR using the Luminaris HiGreen qPCR Master Mix Kit (Thermo Fischer) with a standard amplification protocol (denaturation: 95°C, 15 s; annealing: 60°C, 30 s; 72°C, 30 s) and an equivalent of 5 ng RNA as input. Primer information is given in table 3. Signals were normalized either to RNA Pol II expression levels using the ΔΔCT method.

**table 3.**
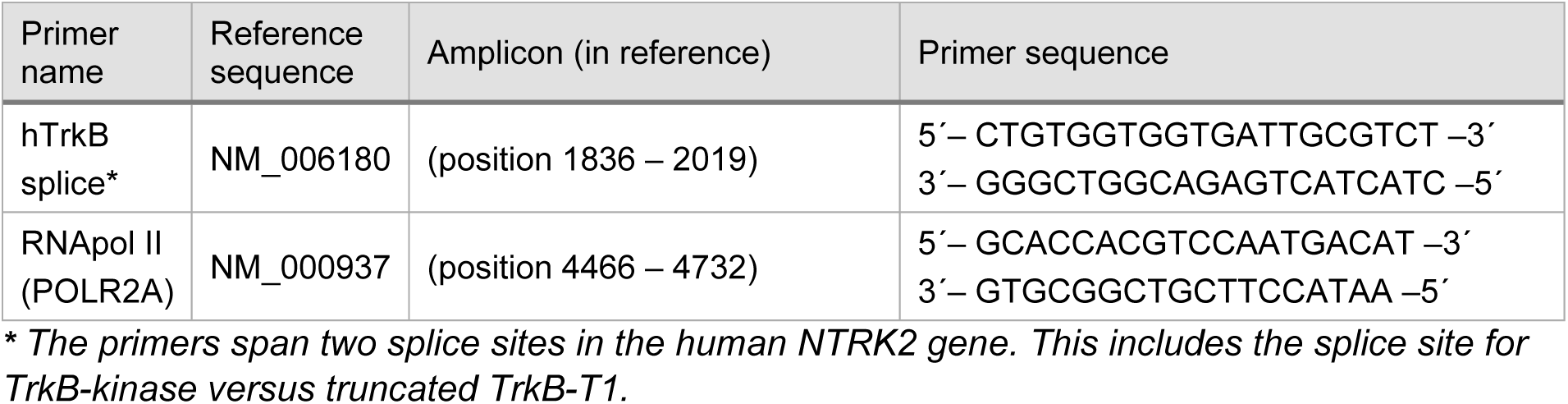
primer sequences.

### Patient glioblastoma tissue

Retrospective investigation of biomarkers in glioblastoma tissue samples with the help of immunohistochemistry and molecular biology methods was approved by our institutional Ethics Committee (#103/14). Glioblastoma tissue was harvested during brain surgery. Fresh tissue was snap frozen in tissue tek (O.C.T compound, Sakura) at −30°C. For fast histological examination, cryosections were labelled with hematoxylin and eosin stain. After histological examination, tissue-tek embedded, tumor-positive tissue was stored at −80°C in a local tissue bank (Institute of Pathology, Neuropathology, University Hospital, Würzburg). Tissue categorized as grade IV glioma was selected and used for Western blotting, reverse transcriptase qPCR or immunohistochemistry. For immunofluorescence labelling, tissue was embedded in paraffin according to standard procedures. For control, paraffin-embedded frontal brain post-mortem tissue was used.

### Immunohistochemistry (glioblastoma tissue)

For immunohistochemical staining of glioblastoma tissue, formalin-fixed paraffin-embedded sections were deparaffinized in 100% xylene and rehydrated using a graded alcohol series (100%, 96%, 70% for 5 min each). For antigen retrieval, specimens were heat-treated for 10 min in 20 mM citric acid buffer (pH 6.0), in a pressure cooker. Sections were rinsed with dH_2_0 and 1× TBS and blocked with a solution containing 10% horse serum, 0.3% Triton X100 in TBS, for 1h at RT. Afterwards, tissues were incubated over night at 4^°^C with primary antibodies. For glioblastoma IHC, the following antibodies were used: anti-human Nestin (R&D Systems, MAB1259, 1:2,400), goat anti-TrkB (R&D Systems, AF1494, 1:1,000), rabbit anti-pY674/675-TrkA (anti-pY706/707– TrkB) C50F3 (1:500). All antibodies were diluted in antibody-dilution buffer (DCS - Innovative Diagnostik Systeme, Hamburg, Germany). Next day sections were washed twice in TBS and treated with the corresponding secondary antibodies (table 2) for 3 h at RT. Sections were washed twice in 1× TBS and stained with DAPI (2mg/ml stock solution, freshly diluted 1:5,000 in 1× TBS) for 5 min. Sections were washed twice in 1× TBS and were finally embedded in Aquapolymount (Polysciences). Positive control for the primary antibody directed against Nestin was kidney tissue. For negative controls, biopsy samples from healthy donors (from autopsies) were used. Cross-reactivity of secondary antibodies was tested by using secondary antibodies in the absence of corresponding primary antibodies.

### Statistical analysis

Statistical analyses were performed with Origin Pro 2019b. Data are presented with standard error of the means (± SEM). Column statistics were run to check for Gaussian distribution to decide whether to use parametric or non-parametric tests. If one of the groups to be analysed failed the normality test or if value number was too small to run the normality test, non-parametric tests were chosen for further analysis. Normality was tested with the Shapiro-Wilk test and Equality of variances (Levene’s test) and based on the results either the 2-sample t-test or the Mann-Whitney-U-test was used. Results were considered statistically significant at p < 0.05.

## Acknowledgements

We thank Michaela Kessler for excellent technical assistance and Svenja Meierjohann and Kurt Bommert, University of Würzburg, for their scientific input. R.G. has been supported by a fellowship of the Graduate School of Life Sciences (GSLS) Würzburg. The work by R.B. has been supported by grants of the Deutsche Forschungsgemeinschaft BL567/3-2 and the Collaborative Research Center TRR58, project A10. This work was further supported by the Deutsche Forschungsgemeinschaft, Collaborative Research Centre SFB TRR225 (B01) to C. V.

## Author contributions

***Rohini Gupta:*** Conceptualization, Methodology, Validation, Formal analysis, Investigation, Writing - original draft, Writing - review & editing, Visualization, Funding acquisition

***Melanie Bauer:*** Investigation

***Gisela Wohlleben:*** Investigation, Methodology

***Vanessa Luzak:*** Investigation, Writing – review & editing

***Vanessa Wegat:*** Investigation

***Dennis Segebarth:*** Formal analysis, Investigation, Writing – review & editing, Visualization

***Elena Bady:*** Investigation

***Georg Langlhofer:*** Investigation

***Britta Wachter:*** Investigation

***Steven Havlicek:*** Investigation

***Patrick Lüningschrör:*** Methodology

***Carmen Villmann:*** Conceptualization, Resources, Writing – review & editing, Funding acquisition

***Bülent Polat:*** Resources

***Camelia Monoranu:*** Conceptualization, Methodology, Resources

***Jochen Kuper:*** Conceptualization, Investigation, Data curation, Writing – original draft

***Robert Blum:*** Conceptualization, Methodology, Validation, Formal analysis, Investigation, Writing - original draft, Writing - review & editing, Visualization, Supervision, Project administration, Funding acquisition

---

**EV1.**
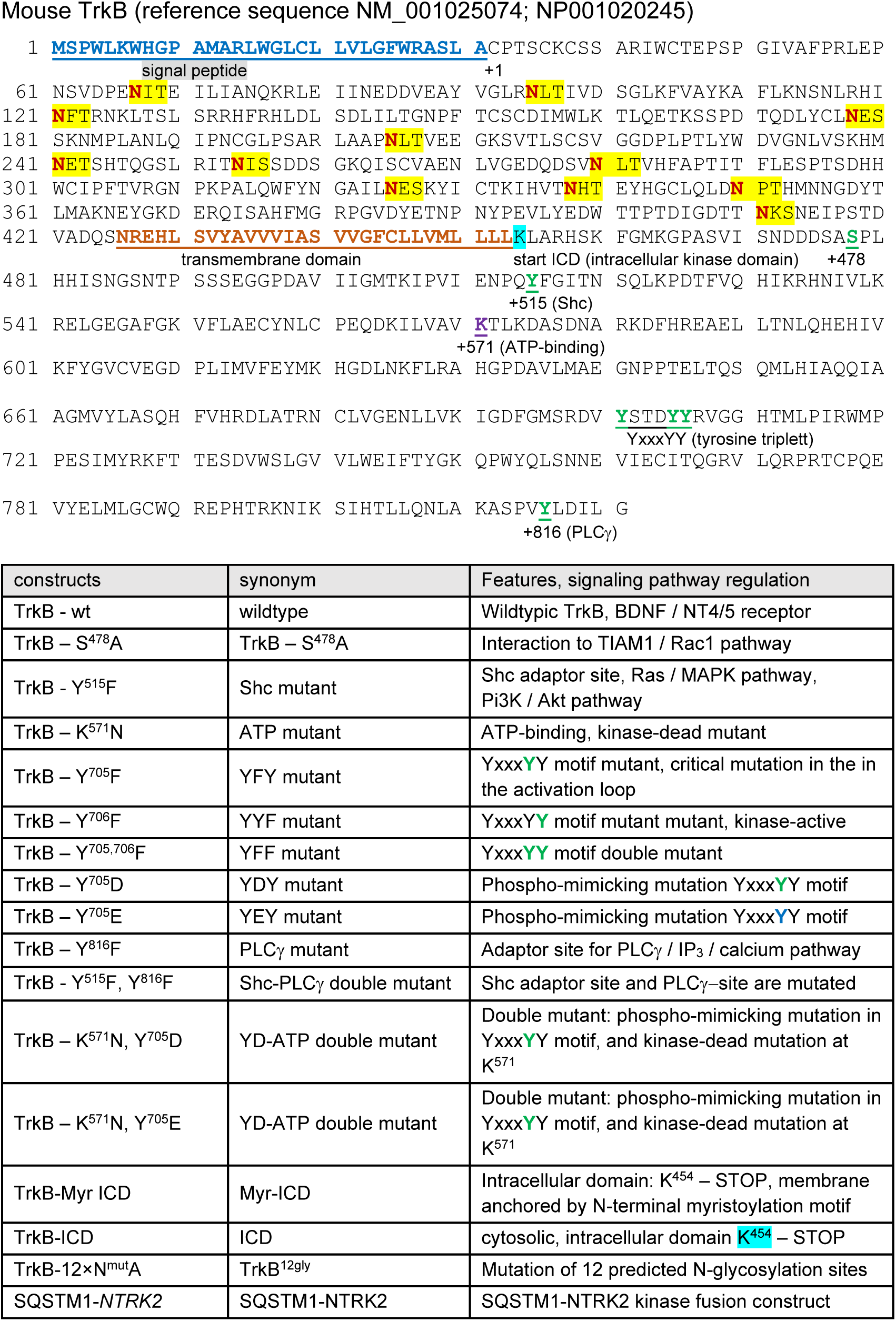
TrkB model and construct. A Deduced amino acid sequence of *Ntrk2* (*trk*B full-length – *trk*B.FL) encoded by *Mus musculus* (reference: NM_001025074; NP_001020245). In the depicted amino acid sequence, in blue is the initiating methionine and signal peptide, in orange is the transmembrane domain, in purple the lysine residue of the ATP binding site and in green are the important serine or tyrosine residues of the kinase domain. Putative N-glycosylation sequons are indicated in red and yellow. B Table explaining the various TrkB constructs generated for use in this study.

**EV2A.**
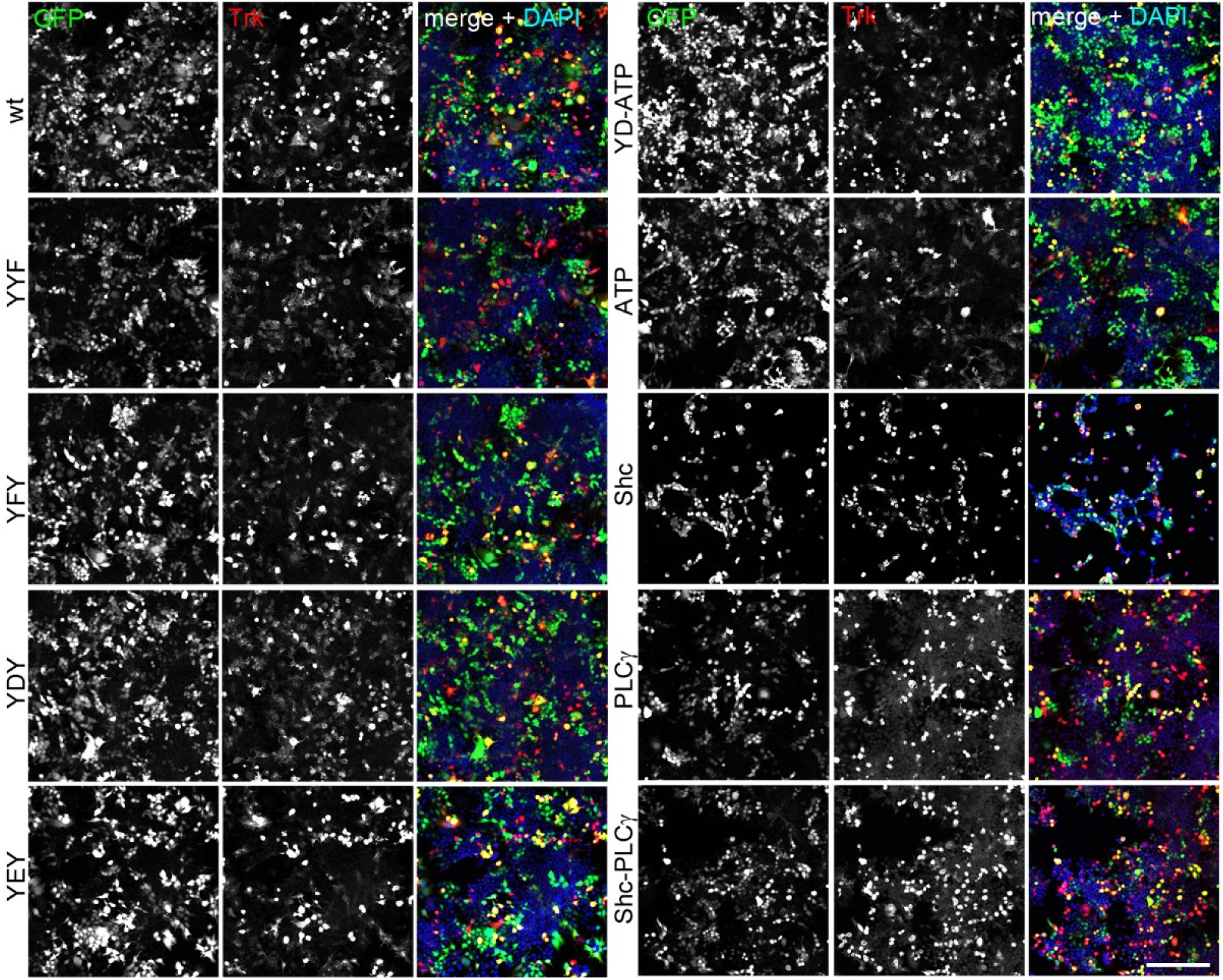
Confirmation of anti-Trk immunoreactivity of TrkB mutants. Immunostaining of HEK293 cells expressing either TrkB wildtype or indicated TrkB mutants. Cells were co-transfected with GFP. Immunofluorescence of GFP (green) and panTrk (red, anti-panTrk (C-term) – A7H6R), together with DAPI as nuclear counterstain (blue). Cells were immunostained after an expression time of 48 h. panTrk binds to the C-terminus of the receptor and shows there the integrity of the open reading frame of all mutants. HEK293 themselves do not express TrkB. Confocal images; scale bar: 100 μm.

**EV2B.**
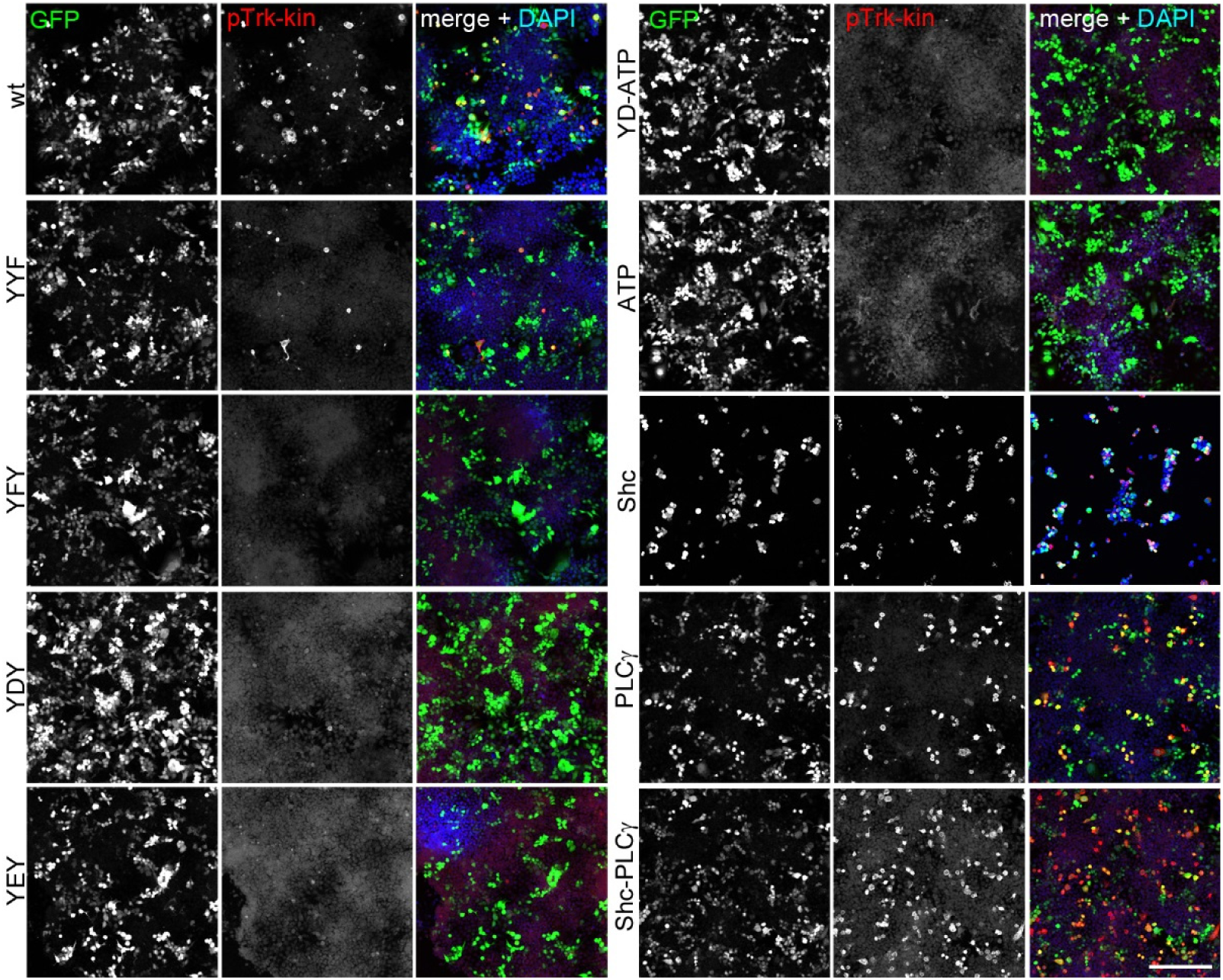
Constitutive activation of TrkB by overexpression and immunoreactivity profile of anti-pTrk-kin. Immunostaining of HEK293 cells expressing either TrkB wildtype or indicated TrkB mutants. Cells were co-transfected with GFP. Immunofluorescence of GFP (green) and pTrk-kin (red, anti-pY674/675-TrkA (anti-pY706/707–TrkB) C50F3), together with DAPI (blue). Cells were immunostained 48 h after transfection. pTrk-kin antibody binds to the phosphorylated 2^nd^ and 3^rd^ Y residues in the YxxxYY motif of the Trk kinase domain. When these sites are mutated (Y^705,706^F), there is no fluorescence seen as the antibody no longer recognizes them (YYF, YFY, YEY). When the ATP site is mutated (K^571^N), lack of ATP prevents TrkB autophosphorylation and the Y residues in the kinase domain remain unphosphorylated (ATP, YD-ATP). All missense mutations in Y705 for D,E,or F, interrupted the anti-pTrk-kin immunoreactivity TrkB. Mutations in the Shc (Y^515^F) and PLCγ (Y^816^F) sites did not affect the phosphorylation of the YxxxYY residues and cells remain positive for pTrk-kin (Shc, PLCγ, Shc-PLCγ). Confocal images; scale bar: 100 μm.

**EV2C.**
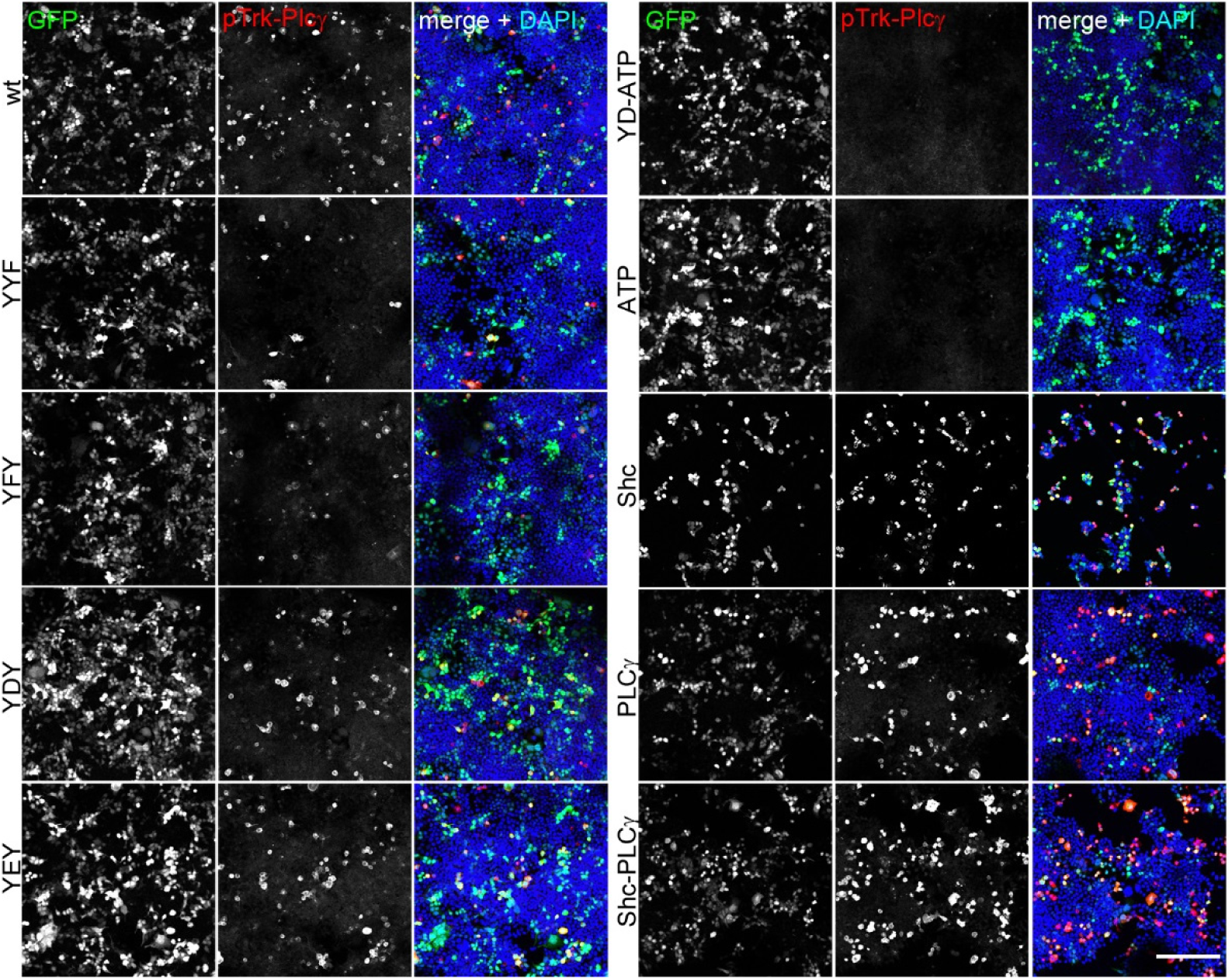
Constitutive activation of TrkB by overexpression and immunoreactivity profile of anti-pTrk-PLCγ. Immunostaining of HEK293 cells expressing either TrkB wildtype or indicated TrkB mutants. Cells were co-transfected with GFP. Immunofluorescence of GFP (green) and pTrk-PLCγ (red, anti-pY785-TrkA (anti-pY816-TrkB) C67C8), and a DAPI counterstain (blue). Cells were immunostained 48 h after transfection. Confocal images; scale bar: 100 μm. In kinase-dead TrkB mutants (ATP, YD-ATP) TrkB remains unphosphorylated. This shows that the antibody is phospho-specific in TrkB. Constitutive phosphorylation is seen in all other mutants, albeit at different intensities. A directed missense mutation at the PLCγ-site (Y^816^F) creates a phospho-independent, immunoreactive site after paraformaldehyde fixation (see also EV2D for anti-pTrk-Shc).

**EV2D.**
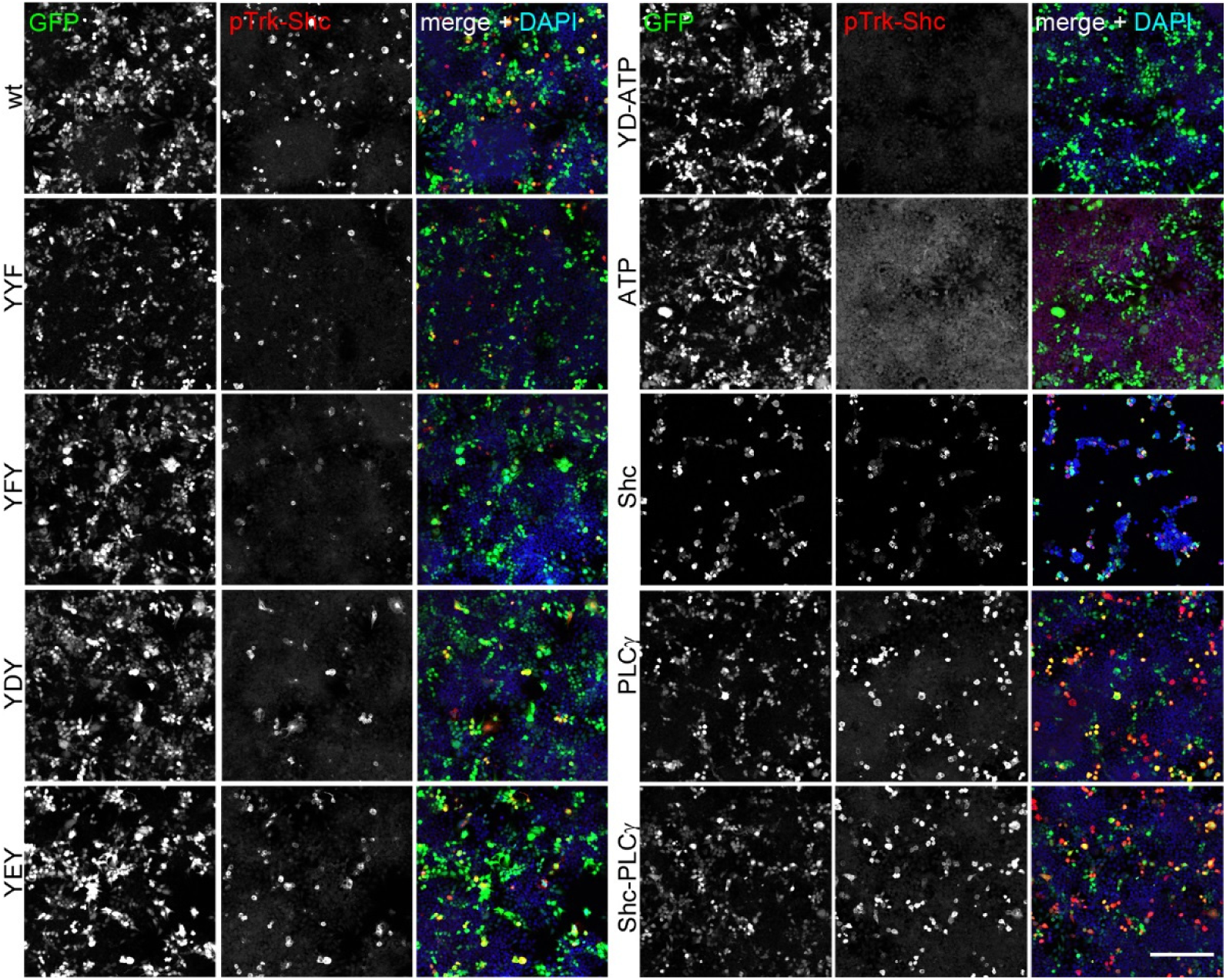
Constitutive activation of TrkB by overexpression and immunoreactivity profile of anti-pTrk-Shc. Immunostaining of HEK293 cells expressing either TrkB wildtype or indicated TrkB mutants. Cells were co-transfected with GFP. Immunofluorescence of GFP (green) and pTrk-Shc (red, anti-pY490-TrkA (anti-pY516-TrkB) C35G9), and a DAPI counterstain (blue). Cells were immunostained 48 h after transfection. Confocal images; scale bar: 100 μm. In kinase-dead TrkB mutants (ATP, YD-ATP) TrkB remains unphosphorylated. This shows that the antibody is phospho-specific in TrkB. Constitutive phosphorylation is seen in all other mutants, albeit at different intensities. A directed missense mutation at the Shc-site (Y^515^F) creates a phospho-independent, immunoreactive site after paraformaldehyde fixation (see Shc and Shc-PLCγ double mutant).

**EV2E.**
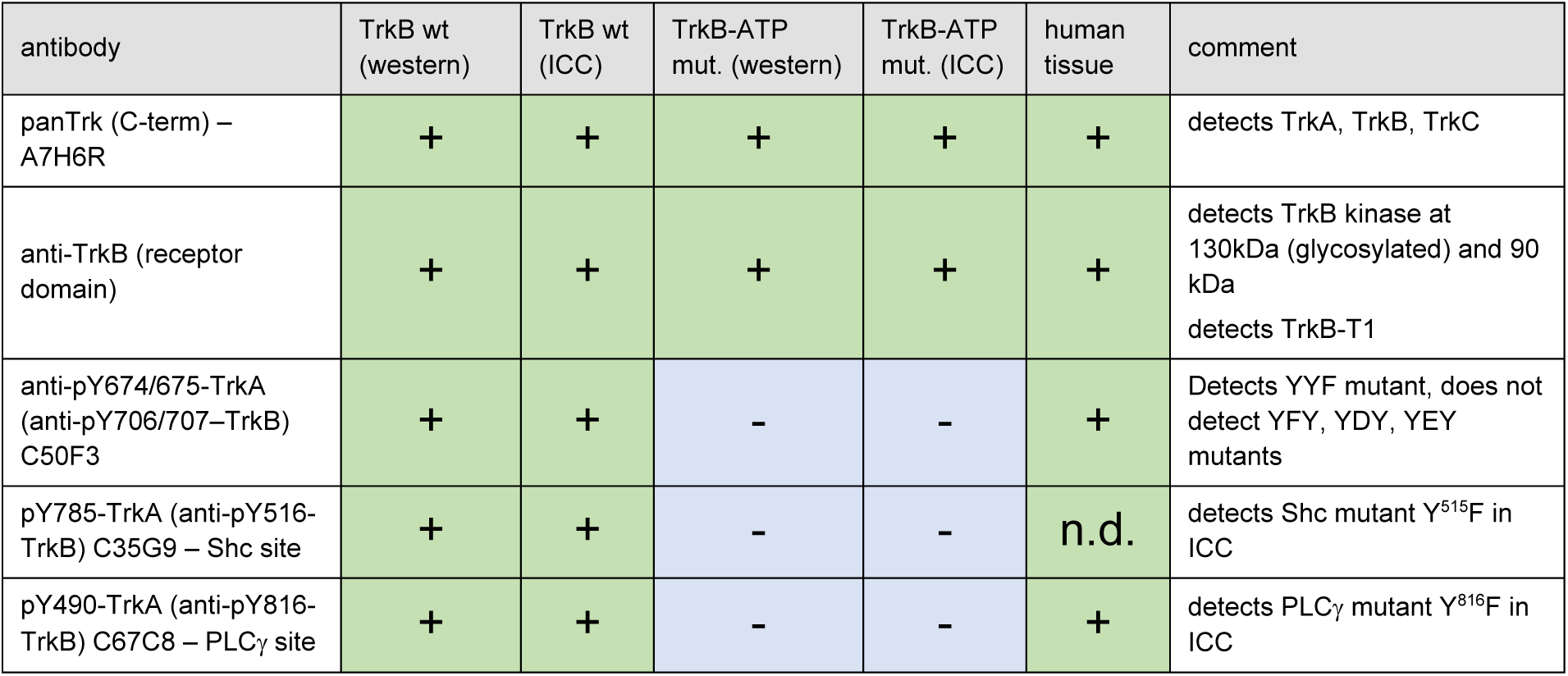
Table describing properties of anti-Trk antibodies used for TrkB detection. In the kinase-dead TrkB-ATP mutant, corresponding tyrosine residues are not mutated, but autophosphorylation is inhibited due to a missense mutation (K^571^N).

**EV3.**
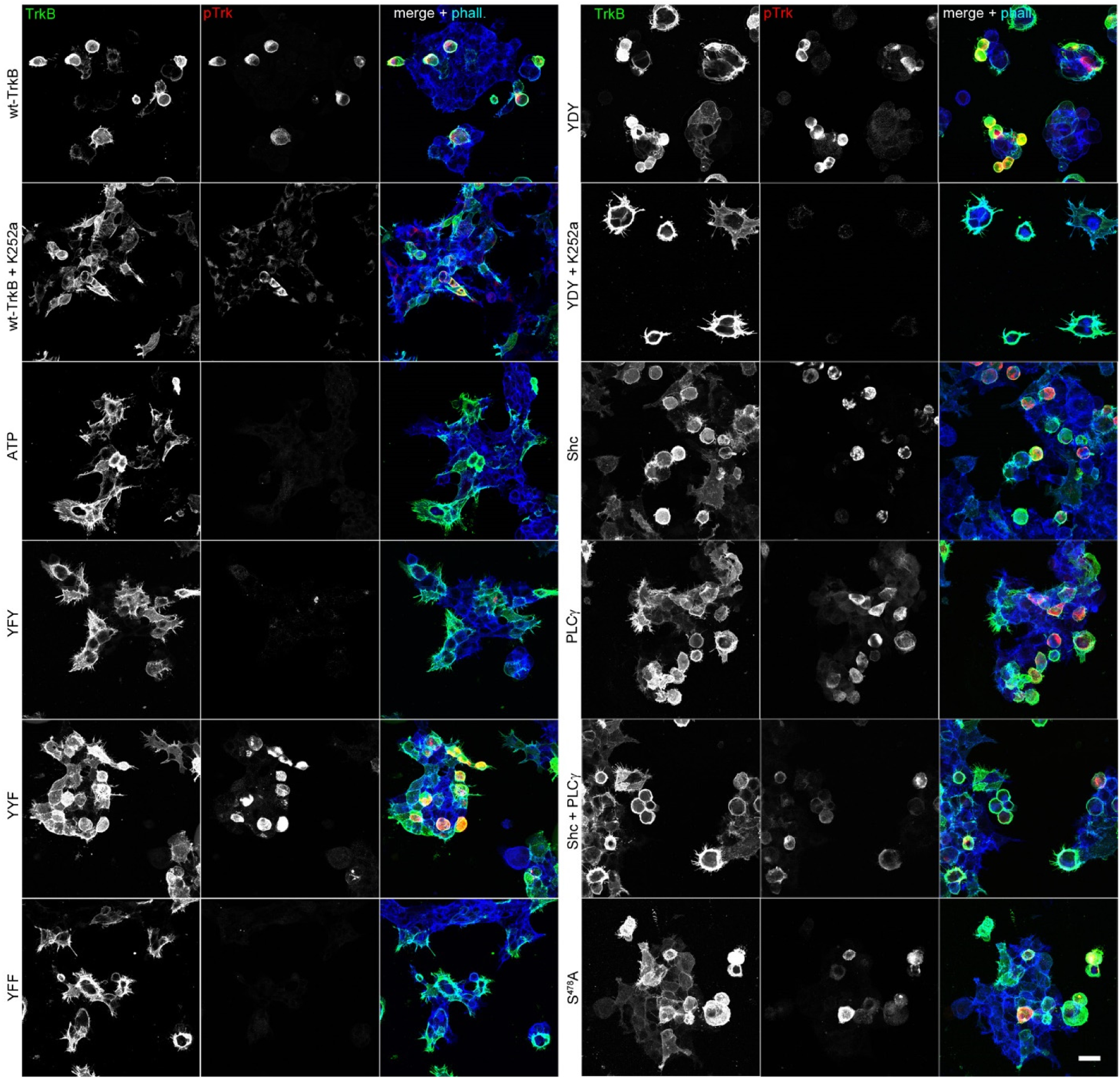
Abundance-dependent auto-activation of TrkB kinase changes in actin morphology (extended visualization for Fig 1 F). Round-shaped cells express kinase-active TrkB mutants. Immunostaining of HEK293 cells expressing either TrkB wildtype or indicated TrkB mutants. Immunofluorescence of TrkB receptor (green) and pTrk-PLCγ (red). PLCγ (Y^816^F) mutants (TrkB-PLCɣ, TrkB-Shc-PLCɣ) were stained with pTrk-kin (red). F-actin was labelled with Acti-stain-670 phalloidin (blue). Typically, filamentous cells express the kinase-dead ATP mutant of TrkB or the YxxxYY mutants (YFY and YFF). Treatment with 150 nM K252a for 30 min reverses the round shape of cells expressing TrkB wt or TrkB-YDY. Confocal images; scale bar: 25 μm.

**EV4.**
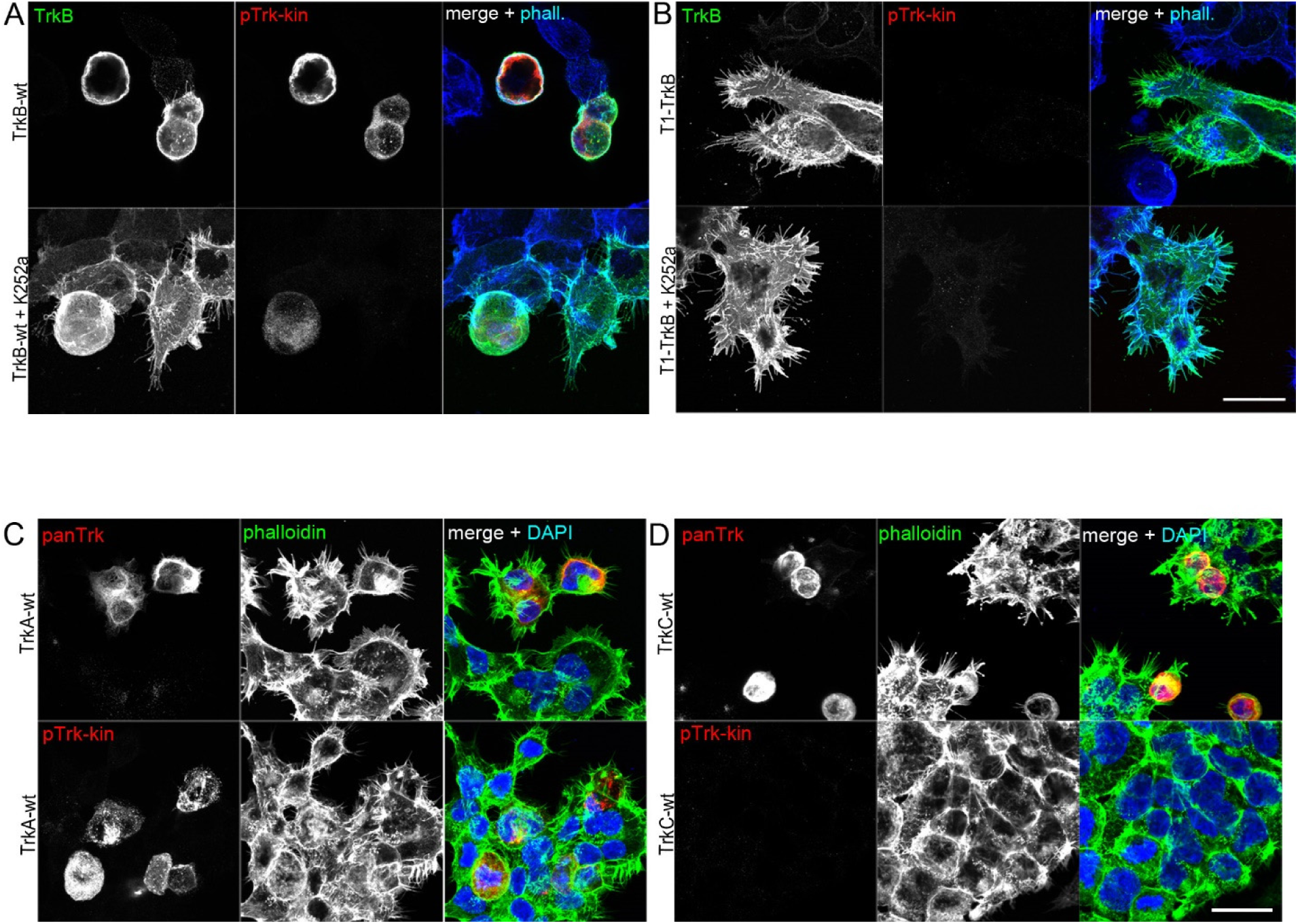
Filopodia formation is preserved in the kinase deficient TrkB-T1 expressing cells, but not in Trk kinase expressing cells. A, B TrkB overexpression causes changes in actin morphology of HEK293 cells. This effect can acutely be reversed by the Trk inhibitor K252a. HEK293 cells expressing the kinase-deficient, truncated slice variant TrkB-T1 remain phospho-inactive and filamentous for control and K252a-treated conditions. Immunofluorescence of TrkB receptor (green) and pTrk-kin (red). F-actin was labelled with Acti-stain-670 phalloidin (blue). Confocal images; scale bar: 25 μm. C, D TrkA and TrkC kinase overexpression leads to roundish cells. Immunofluorescence of panTrk receptor or pTrk-kin (red). Note that TrkC autophosphorylation is not detected by anti-pTrk-kin. F-actin was labelled with Acti-stain-670 phalloidin (blue). Confocal images; scale bar: 25 μm.

## Explanation and detailed description of the results in EV5

Confocal imaging of the TrkB phosphorylation signal suggested that most of the phospho-Trk-wt signal was localized at intracellular sites (Fig 1E). For this reason, we performed a series of experiments to test whether TrkB auto-activation occurs at intracellular sites. First, we expressed TrkB-wt and the kinase-dead TrkB-ATP. Western blotting confirmed TrkB-wt at 90 and 130 kDa. However, after an expression time of about 30 – 48 hours, the TrkB-ATP mutant typically appeared exclusively at 130 kDa (Fig. EV5A), indicating that it was already fully glycosylated. The 90 kDa band of TrkB-wt could be probed with antibodies against the receptor domain (anti-TrkB) and carboxyterminal end of the receptor (anti-pan-Trk) suggesting that the 90 kDa band was not a degradation product. This immature 90 KDa is typically observed after transient expression of TrkB after transfection. In stably expressing TrkB cells, for instance after lentiviral transduction or stable expression, we don’t see this 90KDa TrkB band. Under stable expression, or induced expression from genomic sites, TrkB appears at 130 kDa (see also Fig. 7).

To further test how glycosylation affects the relative molecular weight and constitutive phosphorylation, we mutated first four (TrkB^4gly^), then seven (TrkB^7gly^) and finally 12 (TrkB^12gly^) predicted N-glycosylation sites. Glycosylation sites were predicted with the program NetNGlyc (labelled in EV1), an attempt to distinguish potentially glycosylated sequons from non-glycosylated ones. Mutation of the N-glycosylation sites led to a reduced relative molecular weight and the loss of the 130 kDa band, but the effect of constitutive activation by overexpression remained unchanged, as expected EV5B (see also (Watson et al, 1999)).

For cell surface life labelling of TrkB, we cloned a hemagglutinin-affinity (HA) tag between the signal peptide and the aminoterminal end of TrkB. Putting an HA-tag at this side of the TrkB receptor is known to keep its functionality (Nikoletopoulou et al, 2010). Living HEK293 cells expressing either HA-TrkB-wt or the mutant HA-TrkB-ATP were incubated with an HA-antibody for 15 min at 37°C. Cells were washed to remove residual anti-HA. Fixed cells were permeabilized with Triton X100 and labelled against total TrkB and pTrk-Shc. Secondary antibodies against all three labels were added after permeabilization. High-resolution confocal microscopy showed that cell surface labels against the HA-tag barely overlap with pTrk, thus verifying that most of the pTrk signal was labelled at intracellular sites (EV5C). In cells expressing HA-TrkB-ATP, the cell surface label was also present in filopodia (EV5C) and in vesicle-like structures that are typical after cellular uptake of cell-surface bound antibody/receptor complexes at 37°C (Blum & Lepier, 2008). Vesicle-like pTrk signals were not co-labelled by anti-TrkB (strongly labelled vesicles in the colour-merged image), indicating that these pTrk signals do not carry the epitope for the receptor domain or were not accessible for the label.

**EV5.**
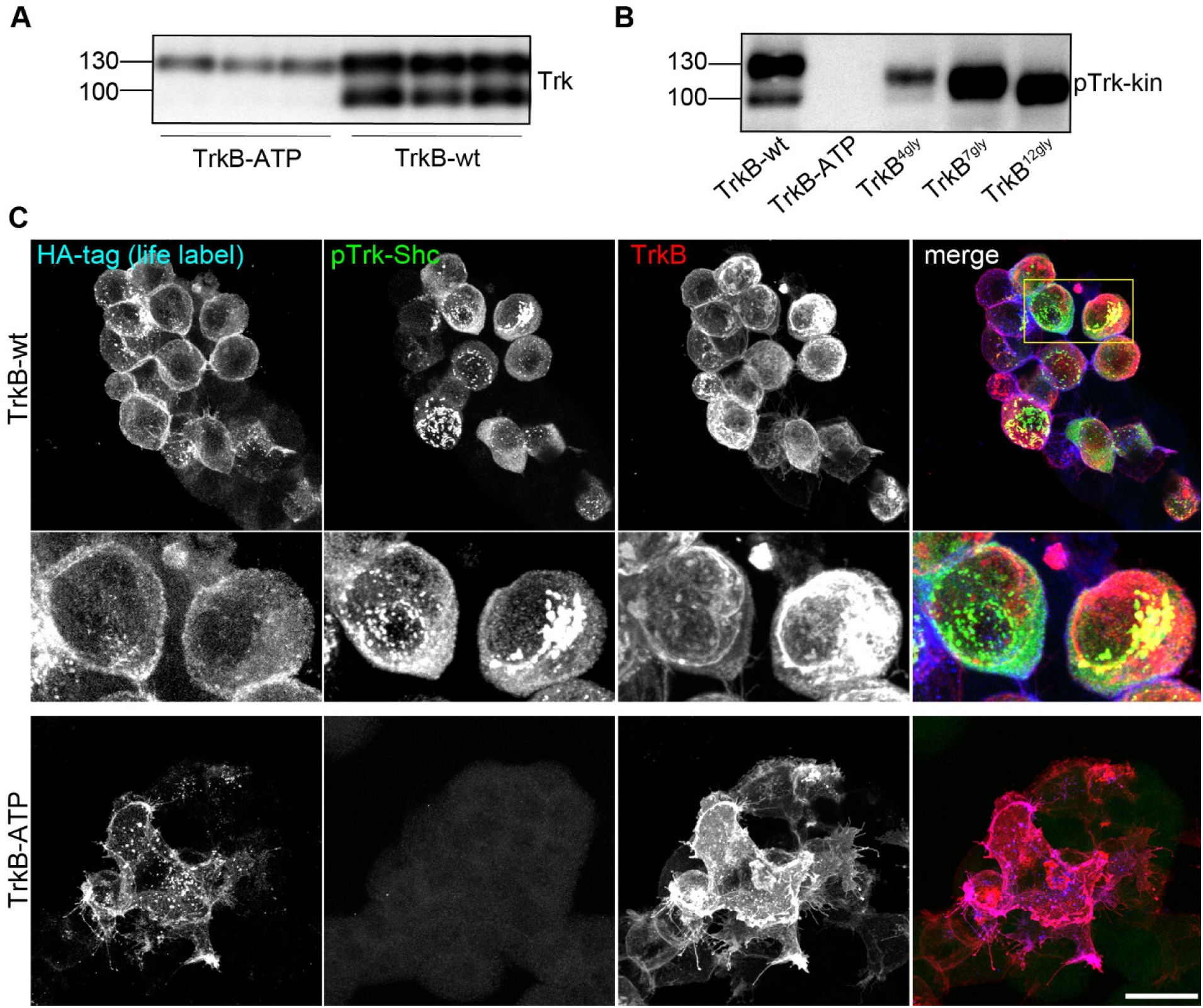
Intracellular localization and delayed glycosylation of constitutive active TrkB. A Western blotting of whole-cell lysates generated from HEK293 cells expressing TrkB-wt or the kinase dead mutant TrkB-ATP. After transient transfection, TrkB was expressed for 30 h. Lysates were probed with anti-panTrk, an antibody that detects TrkB at the intracellular C-terminus. TrkB-ATP shows a Mr of about 130 kDa, while TrkB-wt also runs at 90 kDa, a western blotting band typical for immature TrkB. B Western blotting of whole-cell lysates generated from HEK293 cells expressing TrkB-wt, the kinase dead mutant TrkB-ATP or the TrkB-glycosylation mutants. After transient transfection, TrkB was expressed for 30 h. Lysates were probed with anti-pTrk-kin. All mutants are self-activated except for the kinase dead ATP mutant. The glycosylation mutants appear at different heights depending on the number of mutated glycosylation sites. C Life labelling of TrkB-wt and TrkB-ATP. Cells expressing TrkB were life-labelled via an extracellular HA-tag for 15 min. Then cells were fixed and post-labelled with pTrk and anti-TrkB. Note accumulation of pTrk at intracellular, perinuclear sites and in vesicular clusters.

**EV6. Time-lapse video showing GFP-actin dynamics of HEK293 cells expressing TrkB-wt.** Overview. Time-lapse created from confocal x.y-t image series. Scale bar 25 μm.

**EV7. Time-lapse video showing GFP-actin dynamics of HEK293 cells expressing TrkB-wt.** Detail of membrane blebbing. Time-lapse created from confocal x.y-t image series. Scale bar 25 μm.

**EV8. Time-lapse video showing GFP-actin dynamics of HEK293 cells expressing TrkB-YFF.** Overview. Time-lapse created from confocal x.y-t image series. Scale bar 25 μm.

**EV9. Time-lapse video showing GFP-actin dynamics of HEK293 cells expressing TrkB-YFF.** Detail of filopodia dynamics. Time-lapse created from confocal x.y-t image series. Scale bar 25 μm.

**EV10.**
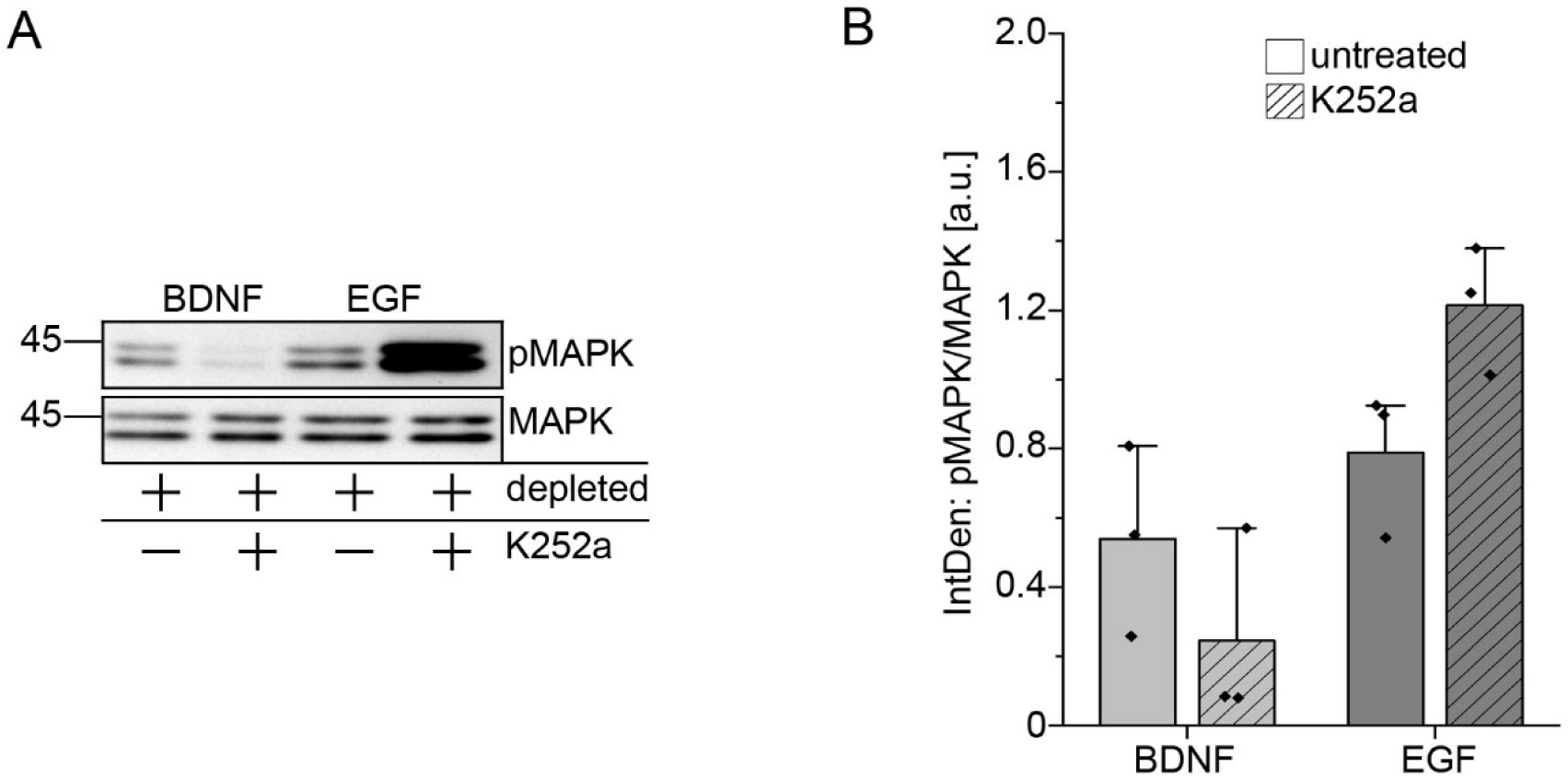
Description of an unexpected off-target effect of K252a. A K252a blocks BDNF-induced MAPK phosphorylation, but potentiates EGF signalling to MAPK. Western blotting experiment with indicated antibodies. Hek293 cells expressed TrkB-wt for 30 h. Serum-depletion (depleted) was performed for 3h. To inhibit TrkB kinase activity, cultures were preincubated with 150 nM K252a for 30 min. DMSO served as solvent control. When indicated, cells were also treated with 20 ng/ml BDNF or 20 ng/ml EGF for 15 min. Western blots show the potentiation of EGF signalling to MAPK. The effect is independent of TrkB and also seen in untransfected HEK293 cells. B Quantification of western blots for pMAPK normalized to MAPK levels with densitometry. Relative integrated densities are shown. K252a treatment for 30 min causes a reduction in TrkB phosphorylation levels after BDNF-stimulation conditions and a potentiation of EGF-induced MAPK phosphorylation. Bar graph: mean ± SEM, overlaid with single data points; n = 3.

**EV11.**
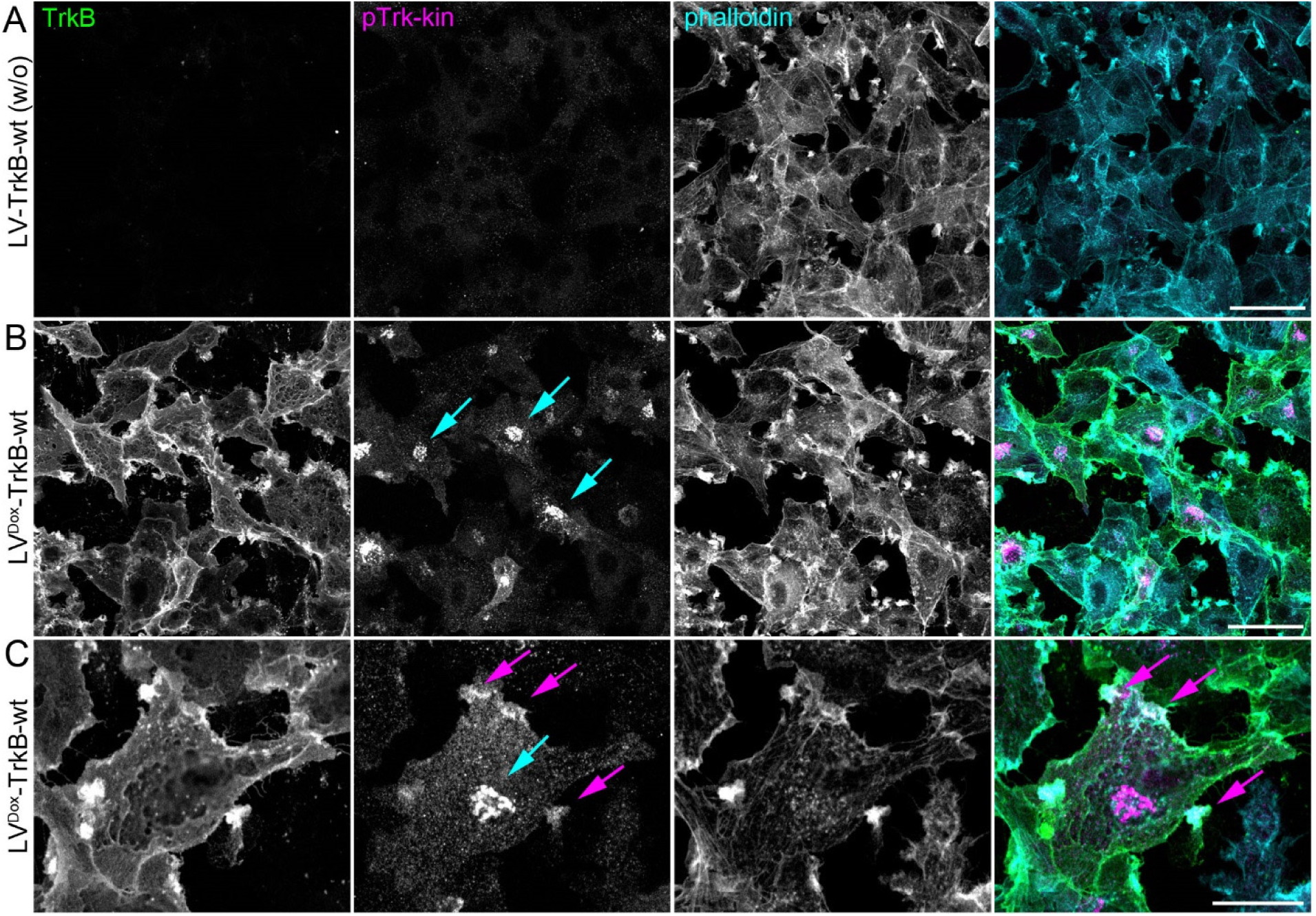
Constitutive active TrkB localizes to the perinuclear Golgi-like region and accumulates in actin-rich protrusions of U87MG cells. Immunofluorescence of TrkB receptor (green) and pTrk-kin (red). F-actin was labelled with Acti-stain-670 phalloidin (blue). A In absence of Doxycyline, TrkB expression is not detectable. Staining as indicated. B Induction of TrkB expression with Doxycycline leads to constitutive activation of TrkB. pTrk signals are pronounced at the perinuclear, Golgi apparatus-like region (cyan arrows). Confocal image; scale bar: 50 μm. C Confocal stack. pTrk-kin localizes also to F-actin rich protrusions (arrows in magenta). Scale bar: 10 μm.

**EV12.**
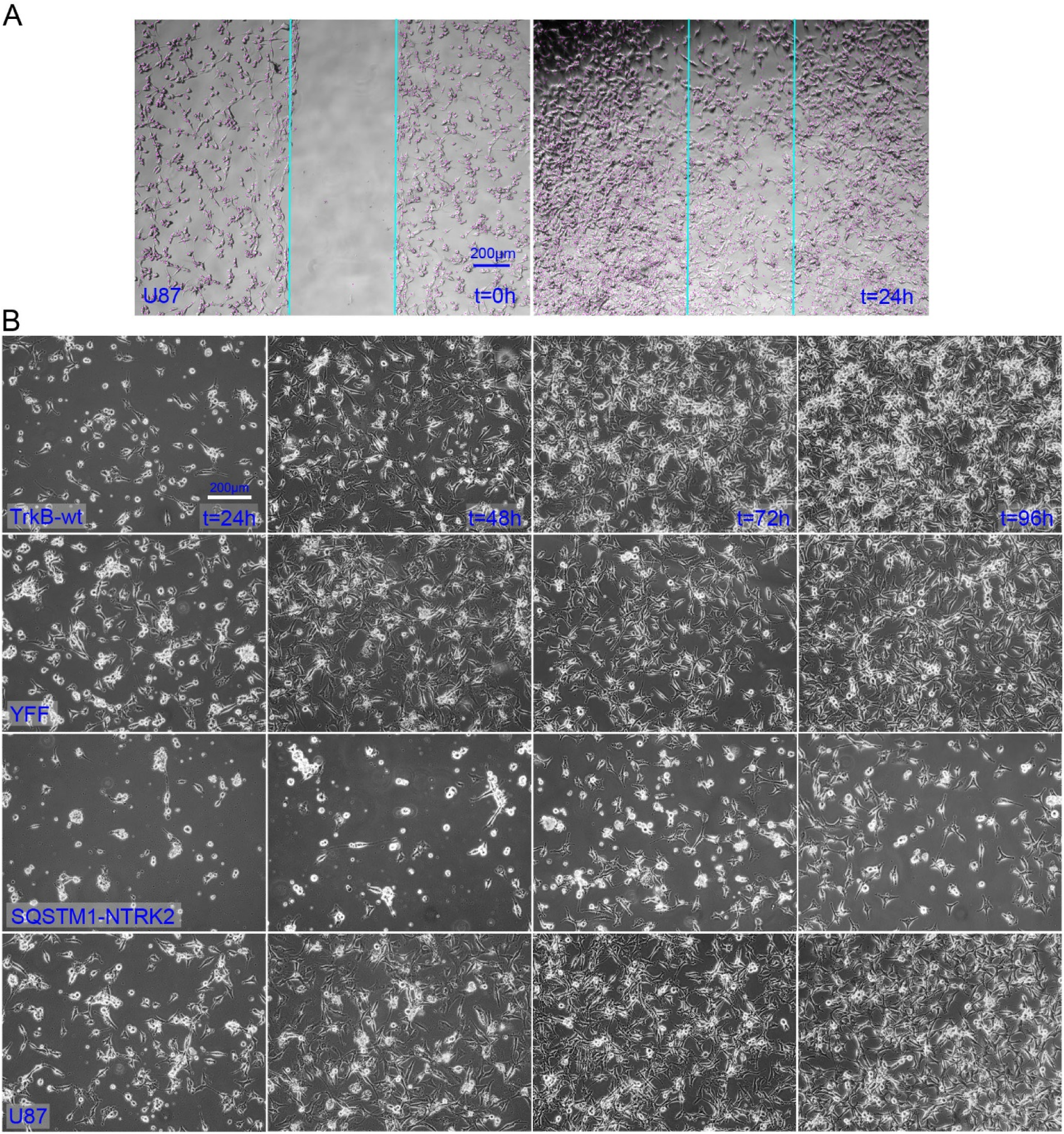
U87MG cells expressing Trk-kinase constructs. A Migration of U87MG cells. Representative phase contrast microscopy images. Indicated in magenta are cells that were automatically counted by unbiased cell counting with ImageJ (see Material and methods). B U87MG cells expressing TrkB-wt or SQSTM1-NTRK2 were not dying within the indicated time span of 96 hours, albeit the cells express a rather high amount of intracellular Trk kinase activity (Fig. 7). Representative phase contrast microscopy images.

**EV13.**
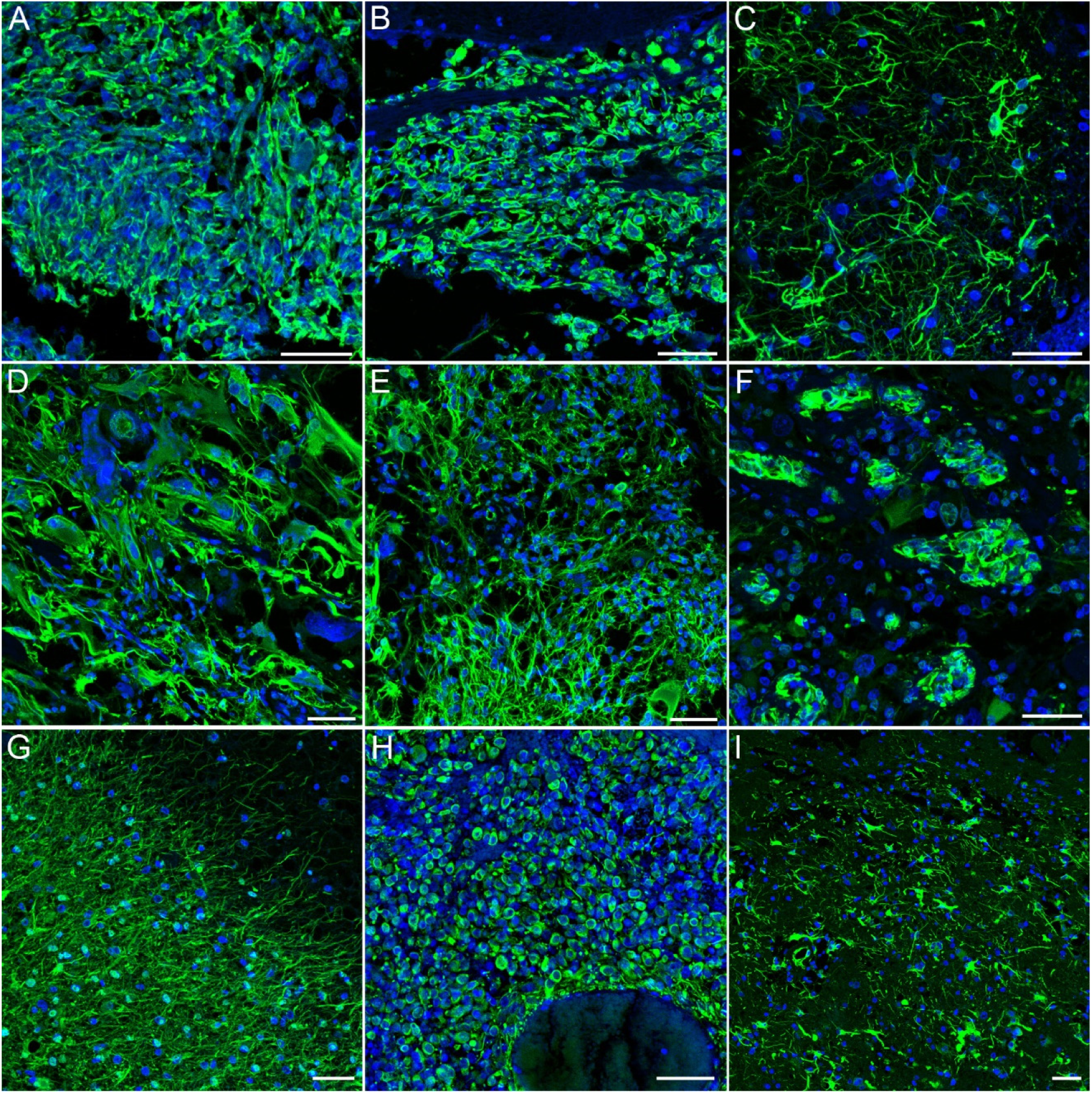
Nestin immunoreactivity in glioblastoma. A-I Nestin (green) immunofluorescence signals in cryosections of post-mortem glioblastoma tissue. DAPI was used as nuclear counter stain (blue). Confocal images, maximum intensity projections, scale bar: 50 μm. In A,B: Different areas of the same patient sample. Nestin+ cells are more roundish and form a globular mass. In C: Neurite-like morphology of Nestin+ cells, In D: Diverse Nestin-morphologies ranging from cells with rather big somata, to smaller cells with disordered neurites. In E: Small spindle-like Nestin+ cells. In F: Individual Nestin+ cell clones. In G: Nestin+ cells ‘stretch’ their neurites in direction of a Nestin-negative area. In H: Small, Nestin-positive cells from a cell mass. The hole in the right lower part of the image shows an intratumoral haemorrhage full of erythrocytes. In I: Nestin+ cells show a disordered neuron-like morphology.

**EV14.**
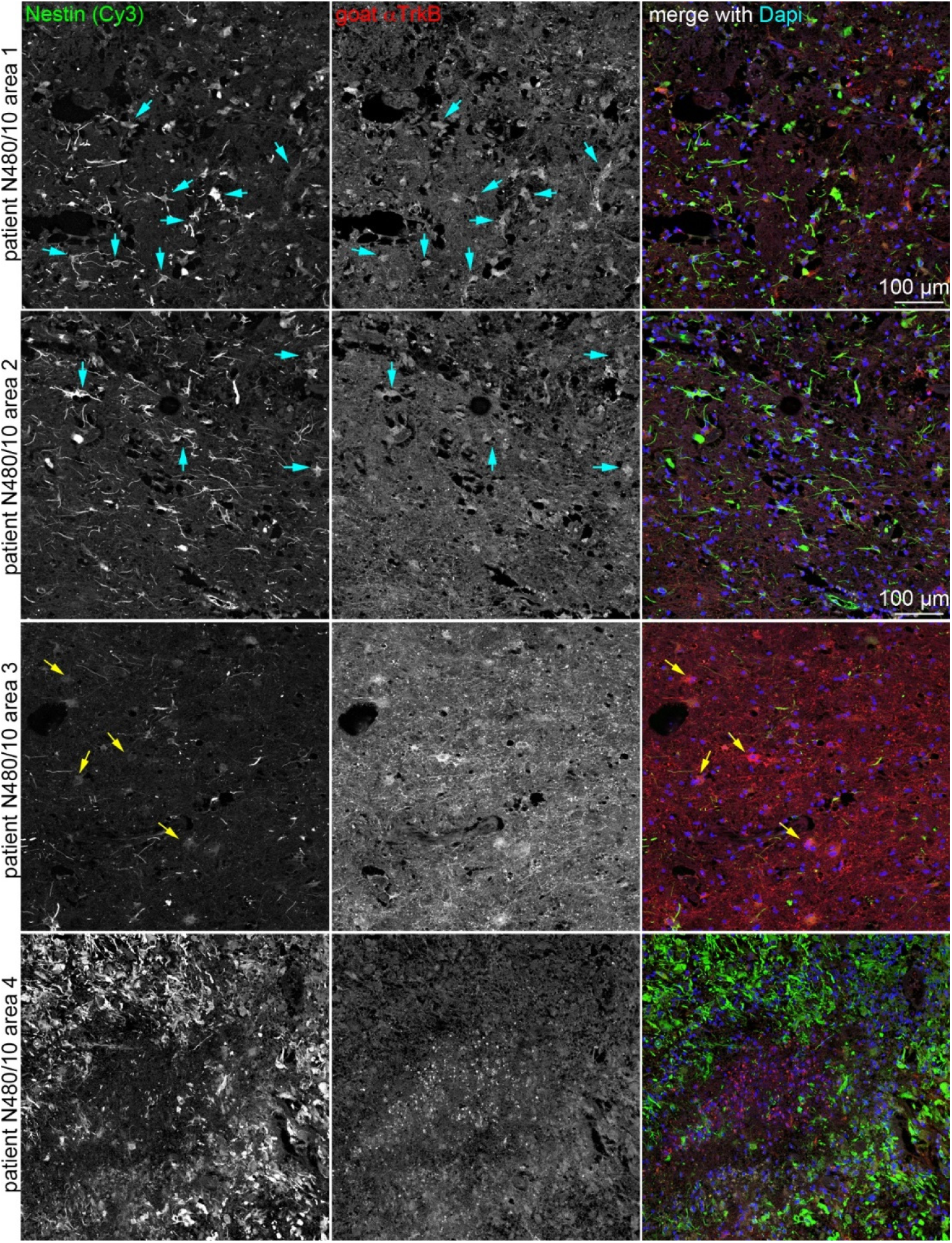
Nestin+ TrkB receptor + cells in glioblastoma and corresponding regional differences in the same sample. Nestin / TrkB positive cells in glioblastoma. Immunofluorescence staining of a representative glioblastoma tissue. Sections were labelled for anti-Nestin, for identification of the glioblastoma and the TrkB receptor. Different areas in the same sample are shown. Confocal laser scanning images. To avoid background fluorescence in the green light channel, Nestin was labelled with Cy3 and TrkB was labelled in far red (Cy5). Dapi was used as counterstain. Many Nestin+ cells show a high abundance of Trk (cyan arrows). In area 3, strong label of TrKB and only rare Nestin immunoreactivity. In area 4, Nestin+ cell masses are organized around a necrosis-like tissue part in the middle of the image.

